# Accurate and fast segmentation of filaments and membranes in micrographs and tomograms with TARDIS

**DOI:** 10.1101/2024.12.19.629196

**Authors:** Robert Kiewisz, Gunar Fabig, Will Conway, Jake Johnston, Victor A. Kostyuchenko, Aaron Tan, Cyril Bařinka, Oliver Clarke, Magdalena Magaj, Hossein Yazdkhasti, Francesca Vallese, Shee-Mei Lok, Stefanie Redemann, Thomas Müller-Reichert, Tristan Bepler

## Abstract

Segmentation of macromolecular structures is the primary bottleneck for studying biomolecules and their organization with electron microscopy in 2D/3D – requiring months of manual effort. Transformer-based Rapid Dimensionless Instance Segmentation (TARDIS) is a deep learning framework that automatically and accurately annotates membranes and filaments. Pre-trained TARDIS models can segment electron tomography (ET) reconstructions from both 3D and 2D electron micrographs of cryo and plastic-embedded samples. Furthermore, by implementing a novel geometric transformer architecture, TARDIS is the only method to provide accurate instance segmentations of these structures. Reducing the annotation time for ET data from months to minutes, we demonstrate segmentation of membranes and filaments in over 13,000 tomograms in the CZII Data Portal. TARDIS thus enables quantitative biophysical analysis at scale for the first time. We show this in application to kinetochore-microtubule attachment and viral-membrane interactions. TARDIS can be extended to new biomolecules and applications and open-source at https://github.com/SMLC-NYSBC/TARDIS.

## Introduction

Electron microscopy^1,2^ (EM) and electron tomography^3–6^ (ET) are leading a revolution in structural biology by enabling high-resolution visualization^7–12^ of complex cellular and macromolecular structures^13–15^. Advancements in EM technology^16^, such as improved microscope optics, detectors, sample preparation techniques, and data collection automation, have significantly increased acquisition speed and quality. As a result, researchers can rapidly collect tomograms and deposit them in databases like EMPIAR^17^ or the CZI data portal^18,19^. Notably, the Electron Microscopy Data Bank (EMDB) has recorded 9,343 tomogram deposits in 2024 alone^20^, and with large-scale imaging initiatives generating thousands of tomograms annually this number continues to grow rapidly.

Despite significant advances and the vast amount of available data, tomogram analysis remains the primary bottleneck, particularly in annotating biomolecules for downstream analyses^21,22^. Accurate annotation is crucial for quantitatively analyzing structures such as microtubules^10,23^, membranes^14^, ribosomes^24,25^, viral particles^13,26^, and other large macromolecules. In contrast to the widely studied particle picking problem in single particle cryoEM/ET^27^, annotation of large biomolecules requires mapping out contiguous objects with complex geometries presenting a unique challenge. However, extracting quantitative insights about these structures requires analyzing numerous tomograms, a task currently limited to only a few datasets due to the time-intensive nature of annotation^7,10,11,22,28–34^. Existing methods aim to accelerate this process, yet annotation can still take days, weeks or even months per dataset due to the lack of streamlined workflows and the need for manual intervention. Large-scale statistical analyses and comparative studies demand tens to hundreds of annotated tomograms, making efficient annotation workflows essential. To fully unlock the potential of tomography for biological research, faster and more automated solutions are urgently needed.

Due to the gravity of the problem, a variety of software tools have been developed for annotating large biomolecules in cryo-EM and cryo-ET data. However, these tools are limited in their generality, accuracy, and speed. Tools either require significant manual effort from users or are limited to specific acquisition methods and biomolecular structures. Furthermore, for large structures, instance segmentation – that is, the problem of separating individual objects of the same type in a tomogram, such as closely packed filaments or membranes – has no acceptable solution. Tools like Fiji^35^, Napari^36^, and IMOD^37^ rely on manual annotation, which, while accurate, is time-consuming and subject to user variability. Others, such as TomoSegMemTV^38^ and MemBrain-seg V2^39^, specialize in membrane segmentation, while Amira^40^ targets microtubules, but these require high-quality tomograms and extensive parameter tuning, limiting their applicability across diverse datasets. General-purpose deep learning tools like Dragonfly^41^ and Micro-SAM^42^ bring automation and interactivity but still demand manual annotations, fine-tuning, or are limited in modality and instance-level segmentation. As a result, researchers often have to rely on multiple tools, investing significant effort to customize each one to suit the specific requirements of individual datasets. No tool seamlessly integrates 2D and 3D instance segmentation across modalities like cryo-ET, cryo-EM, or plastic ET for structures like filaments and membranes in a single, fully autonomous framework without retraining or tuning.

Here, we present TARDIS (Transformer-based Rapid Dimensionless Instance Segmentation)^43^, a modular deep learning framework for annotating biomolecular structures both in EM micrographs and tomograms. TARDIS offers a plug-and-play solution that delivers fast, accurate, and fully automated segmentation across diverse datasets. It supports both semantic and instance segmentation, providing high accuracy instance segmentation for cellular structures for the first time. This is made possible by our state-of-the-art pre-trained networks for membrane and microtubule annotation. To develop these pre-trained models, we created an extensively manually annotated dataset of membranes and microtubules across tomograms and micrographs obtained from both plastic sections and cryo-EM/ET. In total, this dataset contains 382 images with 71,747 objects, the largest manually labeled dataset of its kind to date (**Suppl. Table 1**). As a flexible framework, TARDIS is straightforward to extend with semantic segmentation networks for new macromolecules. These integrate seamlessly with our pre-trained instance segmentation network to automatically identify object instances from the semantic maps, enabled by our novel point cloud-based instance segmentation model architecture, Dimensionless Instance Segmentation Transformer (DIST)^44^, an SO(n) equivariant transformer.

We demonstrate that TARDIS accurately annotates filaments and membranes, improving annotation accuracy by 42% for microtubules and 55% for membranes compared to existing tools. TARDIS is also fast, processing a single tomogram in minutes, orders of magnitude faster than manual annotation, and achieving, on average, a 2x speedup over other algorithms for membrane segmentation and 5x for microtubules. As a pluggable framework, TARDIS can seamlessly integrate with other semantic segmentation tools to provide instance predictions and adapt to new macromolecules and data types. We demonstrate it by applying TARDIS to segment tobacco mosaic virus (TMV)^45,46^ in cryo-ET without any retraining. Moreover, TARDIS streamlined workflow also enables previously infeasible tasks due to computational complexity. We segmented full datasets of a wild-type and two phosphomimetic mitotic spindles in U2OS cells within hours, uncovering variability in kinetochore microtubule (KMT) numbers. In addition, we achieved precise segmentation of liposomes and virus membranes from cryo-ET, thus enabling biophysical characterization of the virus capsids. Moreover, we extended TARDIS by integrating a new semantic segmentation network for actin in cryo-ET and deploying a model for microtubule segmentation in fluorescence microscopy images, all within the same framework. Using TARDIS, we generated the largest high-quality annotation dataset of microtubules and membranes across 13,000 tomograms, a task that would have taken an experienced human annotator approximately 35 years to complete manually. These annotations are now publicly available in the CZI cryo-ET data portal.^18,19^.

TARDIS is open-source and freely available at https://github.com/SMLC-NYSBC/TARDIS, and we also provide an optional Napari plugin-based GUI for TARDIS at https://github.com/SMLC-NYSBC/napari-tardis_em. TARDIS outputs results in all commonly used formats, making them immediately compatible with visualization and analysis tools such as Amira, Napari, and IMOD.

## Results

### TARDIS is a modular framework for any macromolecular segmentation

TARDIS is a framework for fully automated semantic and instance segmentation of cellular macromolecules across diverse resolutions and imaging modalities (**Fig. 1A**). It surpasses existing tools by offering broad applicability without requiring extensive manual parameter tuning. Optimized for accuracy and speed, TARDIS enables rapid annotation of large datasets. Its modular architecture supports seamless extensions to incorporate new macromolecular types (**Fig. 1B**).

**Figure 1:**
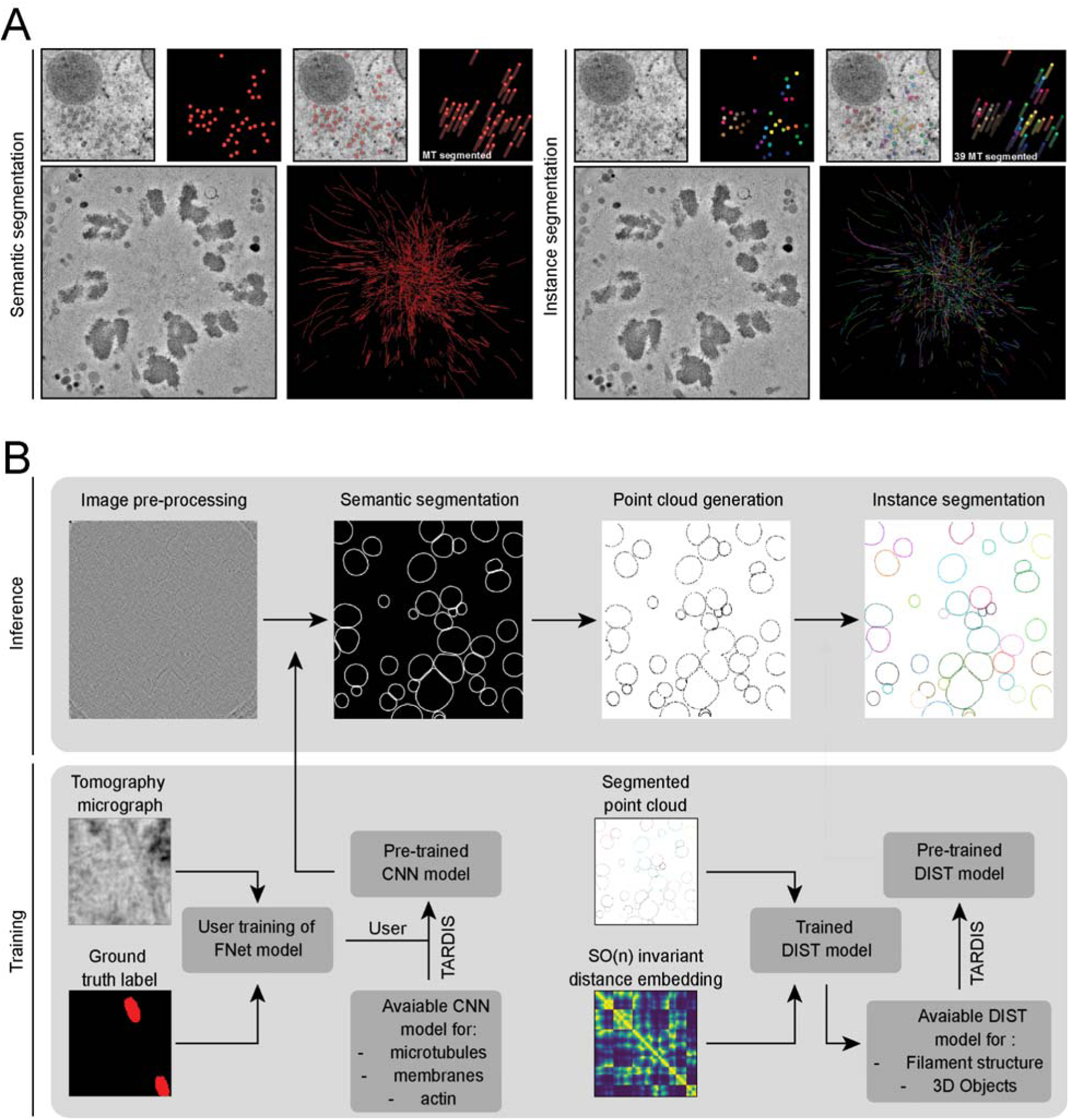
Overview of the TARDIS framework for semantic and instance segmentation. **A)** Example illustrating the distinction between semantic and instance segmentation. **B)** Segmentation of micrographs or tomograms begins with a CNN. TARDIS uses custom pre-trained CNN models to predict actin, membrane or microtubules from tomograms, micrographs and TIRF images. Predicted semantic masks provide the initial segmentation of structures. These masks are then skeletonized to produce a point cloud representing the core structure of each object. Instance segmentation is finalized using the DIST model, which predicts connection probabilities between points in the point cloud. The final instance prediction is obtained through a greedy graph cut.

The framework leverages pre-trained networks for 2D and 3D segmentation tasks and currently includes specialized semantic segmentation models for the following sub-cellular features:

● microtubules in 3D electron tomograms (including cryo and plastic sections)
● microtubules in 2D total internal reflection fluorescence (TIRF) images
● membranes in 3D cryo-electron tomograms
● membranes in 2D cryo-electron micrographs
● actin filaments in 3D cryo-electron tomograms.

By combining these networks with our novel pre-trained point cloud-based instance segmentation model (DIST), TARDIS accurately distinguishes individual structures, enabling quantitative analyses across thousands of tomograms. We provide pre-trained DIST models for two general structures: linear structures (DIST-linear), e.g., filaments or membranes in 2D, and surfaces (DIST-surface), e.g., membranes in 3D. We refer to these collectively as the pre-trained instance segmentation models but apply DIST-linear to filaments and other linear objects and DIST-surface to 3D surfaces when applicable.

### High-Accuracy Semantic Segmentation with FNet

This framework is anchored by our custom-designed FNet, a CNN model for semantic segmentation (**Suppl. Fig. 1**; *Methods*). FNet, a variant of the UNet architecture, provides improved semantic segmentation performance by combining a single UNet-style encoder with two concurrent decoders. One with conventional skip-connections and another with fully connected skip-connections. This architecture improves the precision of cellular structure segmentation, generating probability maps that are converted to binary masks and processed into point clouds (**Suppl. Fig. 2A-B**).

### Geometry-aware instance segmentation with DIST

Point clouds are extracted from voxel-level probability maps through skeletonization, which retrieves the skeletons of segmented structures (**Suppl. Fig. 2A**; *Methods*). The DIST model predicts graph edges between points to delineate individual object instances. Operating on point clouds of any dimensionality, the model utilizes an SO(n) equivariant representation and integrates geometry-aware operations within a transformer network (**Suppl. Fig. 3**; *Methods*). Instances are identified by applying a graph-cut to the edge probability outputs, isolating the most likely connected components as distinct instances.

### TARDIS accurately segments microtubule and membrane instances in tomograms and micrographs

TARDIS aims to alleviate the bottleneck in annotating large tomographic data. To validate its performance, we benchmarked TARDIS against existing tools, such as Amira, MemBrain-seg V2 and TomoSegMemTV, focusing on microtubule and membrane segmentation across datasets of varying quality and resolution (**Suppl. Table 1**). Due to the lack of suitable public data, we created a benchmark dataset comprising hand-curated microtubules and membranes. Specifically, we fully instance-labeled microtubules in 91 tomograms and 112 TIRF images, and membranes in 120 micrographs and 53 tomograms (*Methods*).

Semantic segmentation is evaluated using F1 score, precision, recall, average precision (AP), and precision at 90% recall (P90). We evaluated instance segmentation using the mean coverage (mCov) score, which quantifies the overlap between ground truth and predicted point clouds. These metrics range from 0 to 1, with higher values indicating better performance (*Methods*).

### 3D microtubules segmentation

We evaluated TARDIS on microtubule segmentation task using dataset from both cryo-ET and plastic ET, with resolutions ranging from 16.00 to 25.72 Å per pixel (**Fig. 2A** and **Suppl. Table 1**). Validation results confirmed that the DIST-linear model significantly improves instance segmentation of structures using point clouds, achieving near-perfect scores (**Suppl. Table 2**). Benchmarking on curated datasets further demonstrated TARDIS improvements over Amira, with TARDIS achieving an mCov score of 0.72 compared to Amira 0.42 (**Fig. 2B**). These results underscore TARDIS higher segmentation accuracy, particularly in high-resolution tomograms (≤18 Å), where Amira performance declined (**Fig. 2C** and **Suppl. Fig. 4A**).

**Figure 2:**
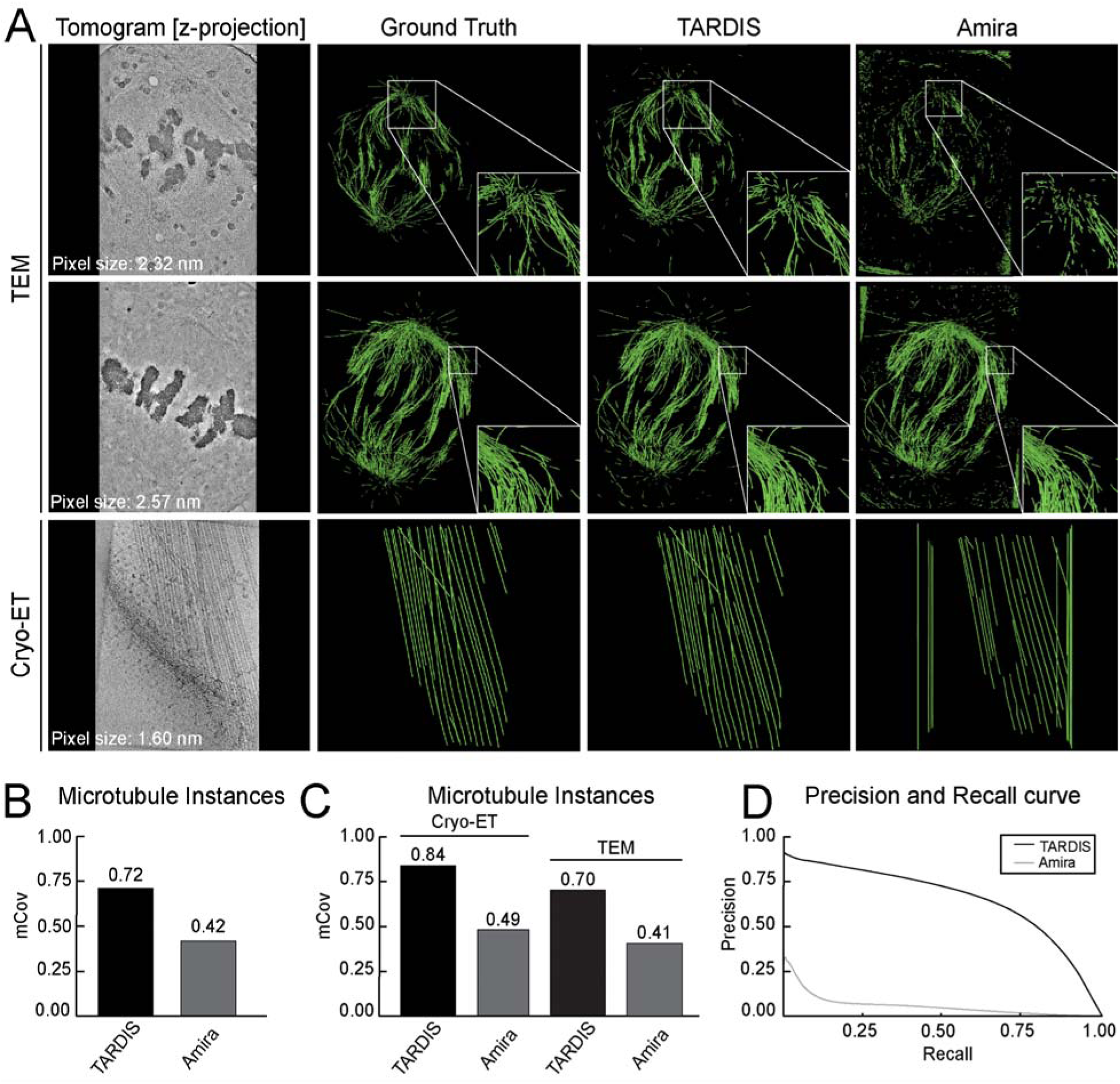
Microtubule segmentation examples and comparison with other state-of-the-art methods. **A)** Instance segmentation examples from TARDIS and Amira, with green lines indicating renders of the core of microtubule filament instances. **B)** mCov metric comparing instance segmentation results from Amira and TARDIS on an entire benchmark dataset. **C)** mCov metric showing instance segmentation results for microtubules using TARDIS and Amira on benchmark datasets from cryo-ET and plastic ET reconstructed volumes. **D)** Precision-recall curves illustrating the quality of semantic segmentation between Amira and TARDIS.

For semantic segmentation, TARDIS consistently achieved a higher F1 score (0.61 vs. 0.17 for Amira), indicating better retrieval of microtubules with significantly fewer false positives. At high recall levels, TARDIS maintained a precision of 0.33, while Amira precision dropped to 0.01 at 90% recall (**Fig. 2D**). Although the Amira template-matching algorithm was previously applied on the plastic ET dataset, its lower accuracy necessitated substantial manual corrections^10,11,28^. In contrast, TARDIS consistently segmented more microtubules with significantly less manual adjustment (**Fig. 2A, Suppl. Fig. 4A and Table 1**).

**Table 1:** Semantic segmentation metrics for microtubule filaments segmented from tomographic dataset. Precision, recall, and F1 scores are calculated using a probability threshold of 0.5 for TARDIS and Amira.

|  | Model | F1 | Precision | Recall | AP | AP90 |
| --- | --- | --- | --- | --- | --- | --- |
| All<br>Datasets | **TARDIS** | **0.61** | **0.70** | **0.55** | **0.69** | 0.33 |
|  | Amira | 0.17 | 0.11 | 0.51 | 0.25 | 0.01 |
| Cryo-ET<br>dataset | **TARDIS** | **0.58** | **0.72** | 0.48 | **0.71** | 0.20 |
|  | Amira | 0.03 | 0.04 | 0.06 | 0.02 | 0.02 |
| TEM<br>dataset | **TARDIS** | 0.61 | 0.70 | **0.56** | **0.67** | 0.35 |
|  | Amira | 0.20 | 0.12 | 0.61 | 0.48 | 0.01 |

### 3D membrane segmentation

We assessed TARDIS for membrane segmentation across varying tomogram resolutions and membrane origins (**Fig. 3A and Suppl. Table 1**). For membrane instance segmentation, we compared TARDIS exclusively with MemBrain-seg V2, which uses a connected component algorithm. TomoSegMemTV was excluded from this analysis due to its frequent generation of single-instance segmentations, making it unsuitable for detailed analyses.

**Figure 3:**
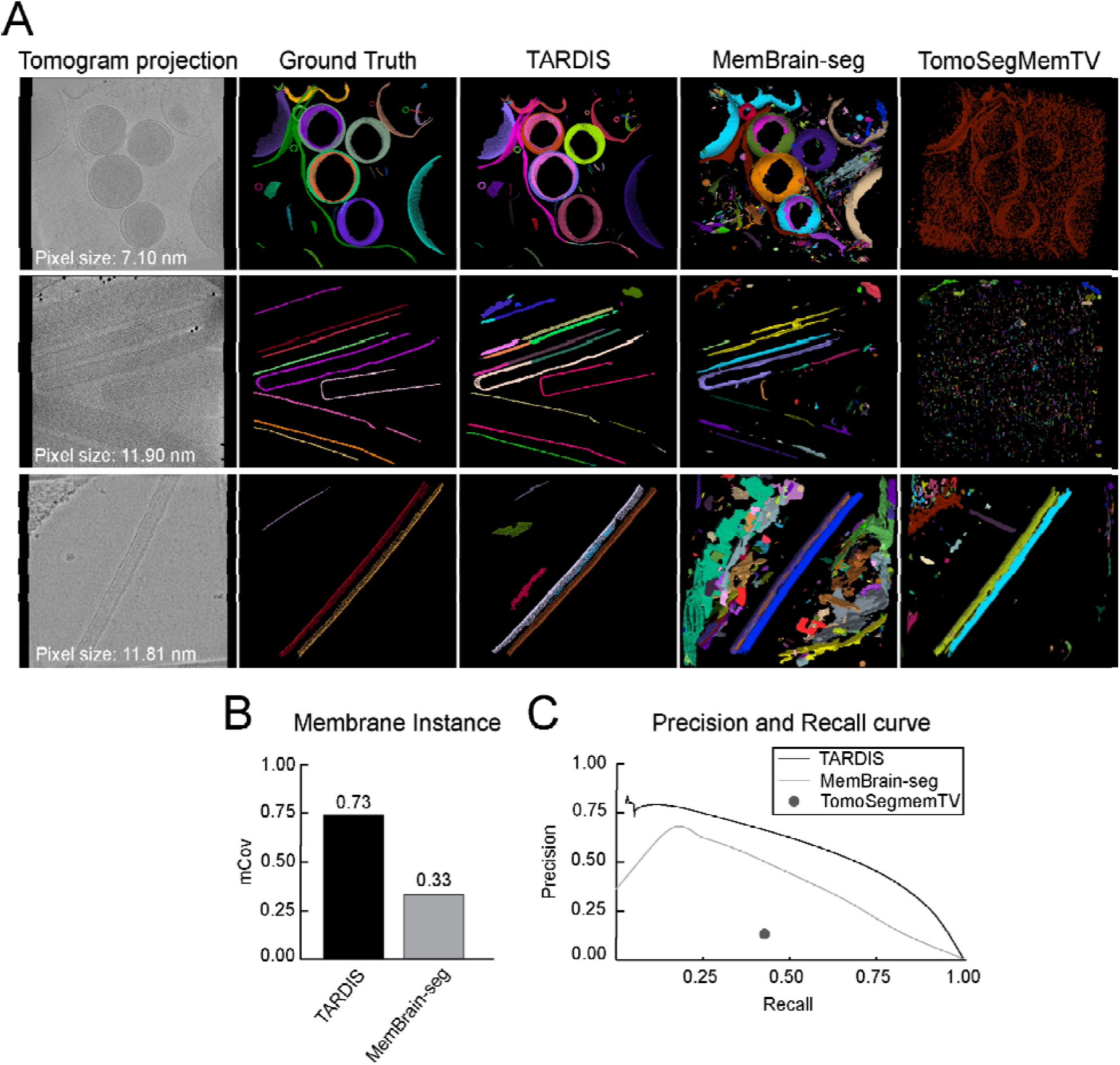
Membrane segmentation from tomographic data examples and comparison with other state-of-the-art methods. **A)** Instance segmentation examples from TARDIS, MemBrain-seg V2, and TomoSegMemTV, with multicolor renderings indicating individual membrane instances. **C)** mCov metric showing a comparison of instance segmentation results between TARDIS and MemBrain-seg V2. TomoSegMemTV was excluded from the comparison as it achieved results close to 0 due to many datasets segmented only as a single instance. **C)** Precision-recall curves illustrating the quality of semantic segmentation between TARDIS, MemBrain-seg V2, and TomoSegMemTV.

We validated that the DIST-surface model improved instance segmentation of surface-like structures using point clouds achieving an almost perfect mCov score of 0.91 (**Suppl. Table 2**). Benchmarking on tomographic datasets further demonstrated TARDIS performance, achieving an mCov score of 0.73, more than doubling the performance of MemBrain-seg V2, which scored 0.33 (**Fig. 3B**). This underscores TARDIS ability to accurately segment membrane instances. Notably, applying the DIST model to membranes initially segmented with MemBrain-seg V2 significantly improved the mCov score from 0.33 to 0.68 (**Suppl. Fig. 6 A-B**).

Beyond instance segmentation, TARDIS significantly improves semantic segmentation through the incorporation of the FNet architecture. We evaluated all methods on semantic segmentation tasks, where TARDIS consistently achieved the highest F1 scores (**Table 2**). Although TARDIS had a lower recall rate of 0.65 compared to 0.72 from MemBrain-seg V2, it demonstrated superior precision-recall trade-offs across various decision thresholds (**Fig. 3C**, **Suppl. Fig. 4B**, and **Table 2**). To further enhance membrane segmentation, we implemented a pixel size normalization strategy. Scaling tomograms to an optimal resolution of 15 Å and micrographs to 4 Å yielded substantial improvements (**Suppl. Results**). Specifically, the FNet model scaled to 4 Å achieved the best results for high-resolution micrographs (<10 Å), while scaling to 8 Å provided the highest accuracy for low-resolution micrographs. Based on these findings, we integrated the optimal pixel size normalization into the automated workflow and publicly released the enhanced model, providing the community with immediate access to this improved tool.

**Table 2:** Semantic segmentation metrics for membranes segmented from cryo-ET.

| Type | F1 | Precision | Recall | AP | AP90 |
| --- | --- | --- | --- | --- | --- |
| **TARDIS** | **0.66** | 0.68 | 0.65 | 0.58 | **0.31** |
| MemBrain-seg V2 | 0.40 | 0.27 | **0.72** | 0.24 | 0.18 |
| TomoSegMemTV | 0.21 | 0.17 | 0.42 | - | - |

### 2D membrane segmentation

We evaluate TARDIS performance on 2D membrane segmentation by training the FNet model for semantic segmentation on a manually annotated dataset. We also incorporated ∼60 nm thick tomogram slices projected into 2D, which we observed to improve model performance (**Suppl. Table 3**). For instance segmentation, we applied the pre-trained DIST-linear model. This approach allowed TARDIS to accurately identify individual membrane instances, achieving the highest mCov score among all tested methods (**Fig. 4A-B**).

**Figure 4:**
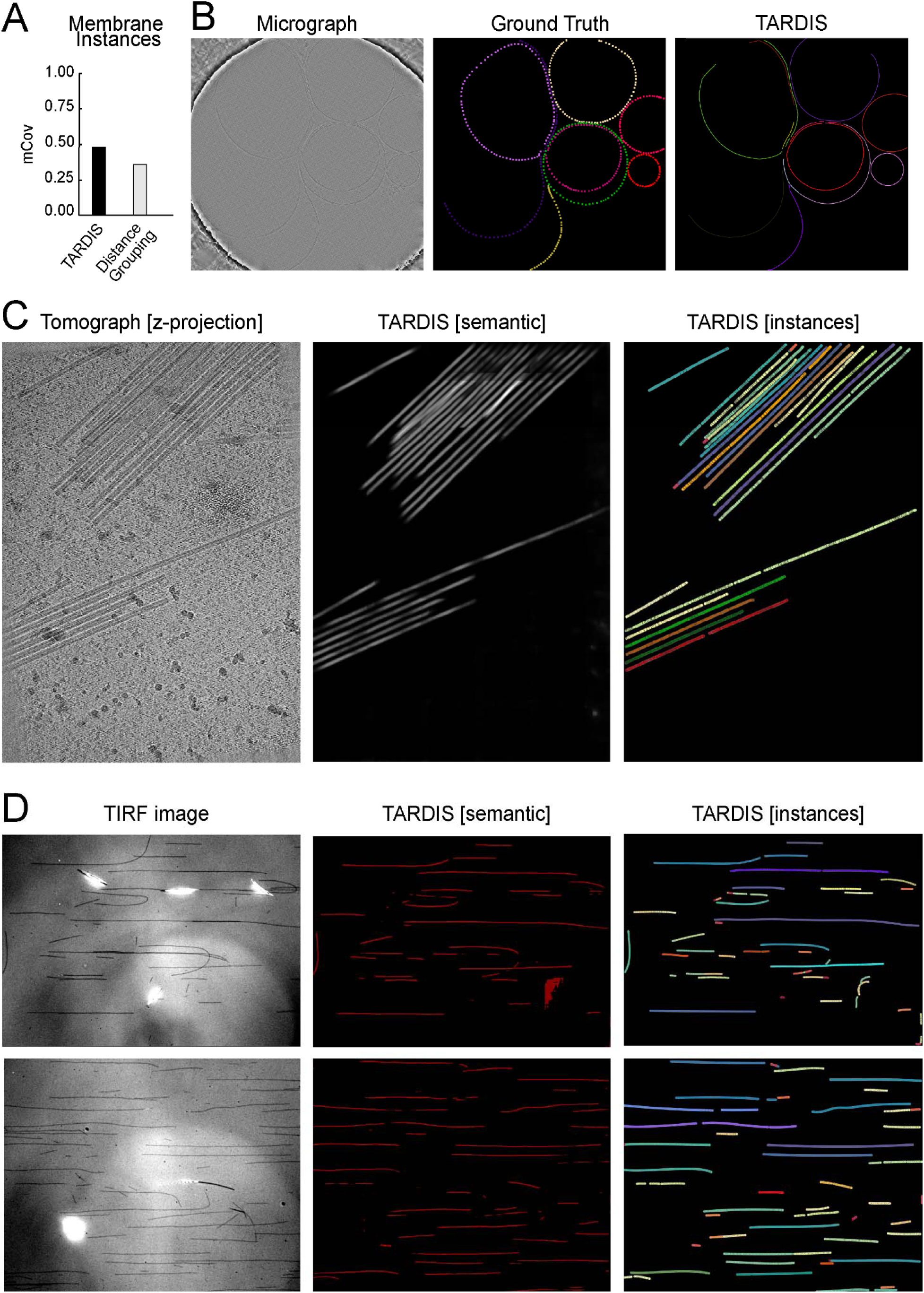
Example of TARDIS applications where one of the models was reused, showcasing TARDIS generalizability. **A)** TARDIS benchmark results on instance segmentation task compared to distance grouping method. **B)** Example of TARDIS instance segmentation of membranes in 2D using the pre-train 2D membrane semantic segmentation model and DIST-linear. **C)** Examples of TARDIS instance and semantic segmentation of Tobacco Mosaic Virus (TMV) filaments, achieved by fully reusing the TARDIS microtubule segmentation workflow. **D)** Examples of TARDIS instance and semantic segmentation of stabilized microtubules from IRM images using TARDIS pre-train FNet model and re-used DIST-linear model.

We tested the pre-trained TARDIS FNet model for semantic segmentation, where it outperformed a similarly deep UNet model, achieving superior F1 scores, particularly in high-resolution micrographs (≤10 Å). TARDIS achieved a precision of 0.74 for membrane structure recovery (**Suppl. Fig. 5A-B; Table 3**). Further analysis demonstrated that the TARDIS FNet model performed well across both high-resolution (≤10 Å) and low-resolution (>10 Å) datasets, achieving an F1 score of 0.41 on lower-resolution images while maintaining high precision (**Suppl. Fig. 5C; Table 3**).

**Table 3:** Semantic segmentation metrics for membranes segmented from cryo-EM. * high-resolution – Micrograph of resolution smaller or equal to 10 Å

|  | Type | F1 | Precision | Recall | AP | AP90 |
| --- | --- | --- | --- | --- | --- | --- |
| All dataset | **TARDIS** | **0.56** | 0.62 | **0.55** | **0.61** | **0.37** |
|  | UNet | 0.54 | **0.64** | 0.51 | 0.61 | 0.37 |
| Dataset with high-resolution* membrane | **TARDIS** | **0.64** | 0.73 | **0.62** | **0.71** | **0.52** |
|  | UNet | 0.62 | **0.74** | 0.58 | 0.71 | 0.52 |
| Dataset with low-resolution** membrane | **TARDIS** | **0.41** | 0.44 | **0.42** | 0.44 | **0.12** |
|  | UNet | 0.40 | **0.47** | 0.37 | **0.45** | 0.10 |

\*\* low-resolution - Micrograph of resolution greater than 10 Å

In summary, the TARDIS FNet model has established a new benchmark for 2D membrane segmentation, demonstrating superior performance across various micrograph resolutions. Moreover, we demonstrated that the TARDIS instance segmentation pipeline can be easily reused. This achievement underscores TARDIS as the first open-source tool capable of reliably segmenting membranes from micrographs and accurately retrieving instances.

### Extending TARDIS to more biological structures and imaging modalities

TARDIS demonstrates high accuracy and efficiency in segmenting microtubules and membranes in electron microscopy datasets, but its versatility extends well beyond these applications. Designed as a modular segmentation tool, TARDIS handles diverse macromolecular structures. To assess this, we evaluated TARDIS on various biological structures and imaging modalities, showing its capacity to generalize across different datasets with minimal or no retraining. We assessed TARDIS ability to accurately segment 2D membranes, actin filaments, and the Tobacco Mosaic Virus (TMV). Given the lack of alternative tools capable of performing instance segmentation for these structures, we identified a distance-based grouping method as a comparison. This method is analogous to the connected component algorithm used by MemBrain-seg V2. All methods were benchmarked using a dataset annotated by an experienced user to assess their effectiveness.

### Segmenting TMV in cryo-ET without TARDIS re-training

To evaluate TARDIS adaptability to different filamentous structures, we applied it to segment the TMV in cryo-ET datasets^47^. For this task, we did not retrain any models. Instead, we employed the pre-trained FNet model, originally developed for 3D microtubule segmentation, alongside the pre-trained DIST-linear model. Despite structural differences between microtubules and TMV, TARDIS successfully segmented TMV filaments without any fine-tuning (**Fig. 4C**). This demonstrates the framework ability to handle diverse filamentous entities with minimal manual intervention.

### TARDIS can segment actin filament trained in limited data scenarios

In many biological studies, limited availability of training data presents a significant challenge for deep learning-based models. To evaluate TARDIS performance under such constraints, we tested it on actin filament segmentation using three annotated tomograms. For comparison, both the TARDIS FNet and a standard UNet model were trained on this small dataset. Despite the limited data, TARDIS outperformed UNet across all metrics (**Suppl. Table 4**) and achieved superior instance recovery, with an mCov score of 0.12 compared to UNet 0.10. These results demonstrate TARDIS robustness and adaptability for novel segmentation tasks, even when training data are scarce.

### Fluorescence microscopy microtubule segmentation

While TARDIS demonstrates effectiveness on EM data, it is designed as a modular tool capable of addressing tasks beyond electron microscopy. To showcase its broader applicability, we tested TARDIS by training the FNet network on TIRF microscopy images of taxol-stabilized microtubules. Utilizing the pre-trained DIST-linear, TARDIS successfully segmented and recovered nearly all microtubule instances from the benchmark dataset (**Fig. 4D**). These highlight TARDIS modular design and its effectiveness in adapting to various imaging modalities while maintaining high performance in instance recovery and segmentation.

### TARDIS enables large-scale annotation for accompanying biophysical analysis and mathematical modeling

In electron microscopy, thousands of tomograms lack segmentation data, as tools are inadequate or the task is deemed too daunting and time-consuming. Yet, instance-segmented datasets are critical for biophysical analysis and mathematical modeling of biological processes, unlocking new knowledge. TARDIS bridges this gap with its speed, accuracy, and fully autonomous operation, segmenting the entire CZI Cryo-ET database in just one month (**Suppl. Results**). We also were able to successfully apply TARDIS on various sample types and obtain high-quality structural data (**Suppl. Results**) in a matter of minutes. This enables novel studies previously out of reach, as they demand vast amounts of annotated data, tasks that once took months or years to complete. It also unlocks complex structural questions, unfeasible before, which require data from diverse conditions with replicates, dramatically scaling up the time needed without such automation.

### TARDIS uncovers variability in KMT number in wild-type *versus* phosphomimetic mutant U2OS cells

During eukaryotic cell division, microtubules connect chromosomes to the mitotic spindle via the kinetochore, a macromolecular complex where the NDC80 complex (NDC80c) serves as the primary microtubule-binding interface^48–50^. The regulation of kinetochore microtubule (KMT) detachment is critical for ensuring accurate chromosome segregation by correcting erroneous attachments^51,52^. Recent biophysical modeling and experimental evidence, including photoconversion and FLIM-FRET assays, have shown that KMT detachment rates are modulated by phosphorylation of the N-terminal tail of Hec1, a subunit of NDC80c^48^. Phosphorylation reduces NDC80c affinity for microtubules, thereby increasing detachment frequency, while dephosphorylation enhances binding stability. To explore this mechanism ultrastructurally, we utilized U2OS cell lines expressing wild-type (WT) Hec1, as well as 9A (all nine phosphosites mutated to alanine, mimicking dephosphorylation) and 9D (all phosphosites mutated to aspartic acid, mimicking constitutive phosphorylation) mutants. Previous studies demonstrated that KMT lifetime decreases with increased phosphorylation (e.g., 9A –> 9D). However, the ultrastructural consequences of these biochemical changes on KMT organization and kinetochore binding remained unclear.^48^.

Given the dynamic nature of KMT attachments, a statistically robust analysis requires imaging and segmenting a large number of cells to account for variability. Traditional manual segmentation or semi-automated tools like Amira are impractical for such scales, often requiring years to process sufficient datasets. To demonstrate the feasibility of such ambitious ultrastructural studies, we employed TARDIS, a fully automated segmentation tool, to analyze three cells, one from each condition, as a proof of concept (**Fig. 6A-C**).

TARDIS analysis revealed striking differences in KMT organization across these conditions. In full-spindle reconstructions, TARDIS segmented 14,452 (WT), 8804 (9D), and 10,366 (9A) microtubules (**Fig. 6D-F**), with 1,228 (WT), 61 (9D), and 251 (9A) identified manually by an experienced researcher as KMTs (**Fig. 6G-I**). The number of KMTs per kinetochore averaged 10.2 ±2.5 (WT), 1.58 ±0.75 (9D), and 6.5 ±1.8 (9A) (**Fig. 6J**). Additionally, we quantified microtubule length distributions, with total microtubule lengths of 10.47 ±9.68 (WT), 8.80 ±5.83 (9D), 9.86 ±7.97 (9A) μm and KMT lengths of 3.22 ±1.43 (WT), 3.39 ±1.74 (9D), 1.23 ±0.77 μm (9A) (mean ±standard deviation; **Fig. 6K**).

These ultrastructural data align with previous observation that phosphorylation of Hec1 tail (mimicked by 9D) reduces NDC80c affinity for KMTs, leading to a shorter KMT lifetime (2.6 ± 0.4 min in 9D vs. 4.0 ± 0.4 min in 9A). The drastically reduced KMT count in the 9D mutant (61 vs. 1,228 in WT) suggests that phosphorylation destabilizes KMT attachments to the point of near-complete detachment. Conversely, the higher KMT count in 9A (251 vs. 61 in 9D) supports the idea that dephosphorylation enhances NDC80c binding affinity, by stabilizing KMT attachments. These findings may provide a structural basis for the nonlinear regulation of KMT stability by Hec1 phosphorylation, as reported in the study, and highlight how biochemical changes translate into physical rearrangements at the kinetochore-microtubule interface.

However, these observations are based on only one cell per condition. The variability in KMT numbers and lengths could reflect cell-specific differences or poor expression of the phosphomutants rather than consistent effects of phosphorylation state. Expanding this analysis to dozens of cells per condition is essential to confirm these patterns. TARDIS makes this feasible by reducing segmentation time from months to hours, e.g., ∼50 hours for three cells versus ∼3 months with Amira (based on an 8-hour workday), a 40-fold improvement. Unlike Amira, which demands labor-intensive manual tracing and refinement, TARDIS autonomously processes large datasets, enabling high-throughput studies critical for mathematical modeling and biophysical analyses. Scaling such an approach for statistical robustness or additional mutant comparisons would exponentially increase this burden, rendering large-scale ultrastructural studies impractical. These limitations underscore the critical role of TARDIS, which delivers accurate, high-throughput segmentation of large datasets, facilitating novel insights into KMT dynamics within feasible timeframes.

### TARDIS identifies virus-liposome interactions quantification

The initial stages of viral infection hinge on precise structural interactions between the virus and host cell^53^. For enveloped viruses like Dengue fever virus (DENV), infection begins with attachment to cell surface receptors, triggering receptor-mediated endocytosis^54^. Once internalized within endosomes, the acidic environment (pH ∼5–6) induces conformational changes *in viral* envelope proteins, facilitating fusion between the viral and endosomal membranes^55^. Understanding the structural dynamics of viral envelope proteins during endosomal fusion is critical for elucidating infection mechanisms and designing antiviral strategies. To investigate these changes, we employed an *in vitro* system using DENV and liposomes as a model for virus-endosome interactions, leveraging high-resolution ultrastructural analysis to capture membrane fusion intermediates.

We utilized TARDIS to analyze DENV-liposome interactions (**Fig. 7A-C**), capitalizing on its ability to perform fully automated segmentation of virus and liposome membranes from electron micrographs without requiring prior training. TARDIS identified interaction sites by segmenting DENV and liposome membranes, representing them as point coordinates paired with segmentation masks (**Fig. 7A**). Object sizes were estimated by fitting spheres to each segmented instance, with DENV classified as objects ∼50 nm in diameter^56^ and larger objects designated as liposomes (**Fig. 7C**). Interaction sites were defined using a 30 nm threshold from the DENV center, reflecting the approximate reach of viral envelope proteins during fusion (**Fig. 7B**). This automated pipeline significantly enhances throughput and precision over manual segmentation, enabling structural analysis of large, heterogeneous datasets *in situ* and providing quantitative insights into DENV-liposome fusion dynamics.

In summary, TARDIS represents a transformative advance in ultrastructural biology by delivering rapid, accurate, and fully automated segmentation, eliminating the need for labor-intensive manual processing. This efficiency overcomes a longstanding bottleneck in structural biology, enabling researchers to generate extensive, high-quality datasets (such as full-spindle reconstructions) critical for mechanistic studies, comparative analyses and biophysical modeling. In the context of DENV-liposome interactions, TARDIS enables detailed quantification of fusion intermediates, offering a scalable approach to probe viral entry mechanisms.

## Discussion

Structural biology has made remarkable strides with advancements in imaging and computational methods, yet tomogram segmentation remains a bottleneck due to the complexity of biological structures. TARDIS addresses this challenge as a novel, plug-and-play tool that combines state-of-the-art semantic and instance segmentation capabilities, offering unmatched precision and efficiency. Unlike traditional tools such as Amira or MemBrain-seg V2, TARDIS eliminates the need for retraining or manual parameter tuning, making it adaptable to diverse datasets and imaging modalities.

TARDIS addresses this challenge by offering a robust, automated workflow for instance segmentation, enabling accurate analysis of complex biological structures, such as microtubules, actin filaments and membranes, where macromolecules coexist in close proximity. A key feature that sets TARDIS apart is its flexibility. Unlike other tools that require retraining for each new segmentation task, TARDIS adapts seamlessly to diverse cellular architectures. This unique generalization capability ensures that TARDIS remains highly relevant as a versatile tool for structural biologists working with evolving datasets and workflows, making it indispensable for modern research.

At the core of TARDIS is FNet, a specialized convolutional neural network architecture designed to enhance semantic segmentation. Our results show that TARDIS outperforms traditional methods, such as UNet, by generating sharper probability masks and capturing finer structural details (**Suppl. Fig. 9A-C**). Additionally, TARDIS integrates the DIST model to identify individual macromolecule instances from the semantic masks produced by FNet. This effectively solves issues of over-segmentation and under-segmentation, which are common pitfalls in other segmentation tools. The DIST model also showcases its broad applicability by seamlessly handling various cellular macromolecules, and even extending to point cloud data, such as LiDAR scans (**Supp. Results** and **Suppl. Fig. 9D**). This highlights TARDIS unique ability to generalize across different types of biological and environmental data.

A major advantage of TARDIS is its fully automated approach. It operates effectively across various dataset modalities, resolutions, and quality levels without requiring hyperparameter tuning or model retraining. This versatility streamlines the segmentation process and minimizes the need for manual intervention, making TARDIS an invaluable tool for structural biologists. To further enhance user interaction, our Napari plugin enables real-time observation of segmentation results and facilitates a more intuitive user experience.

Early adoption of TARDIS within the research community has already demonstrated its practical utility^26,57^. Researchers have successfully applied^19,57–61^ TARDIS to various biological studies, confirming its ability to deliver high-quality segmentations across a range of complex structures. These early use cases have established TARDIS as a key resource for structural biologists and have driven its widespread adoption.

TARDIS represents a paradigm shift in structural biology, offering a precise, flexible, and fully automated solution for tomogram segmentation for numerous different tasks. As an example, with TARDIS it is now possible to systematically analyze wild-type and mutant spindles in mitotic mammalian cells; a comparison that could not be achieved so far within a reasonable time frame. By leveraging the advanced FNet architecture and the integrated DIST model, TARDIS surpasses the capabilities of existing tools, such as UNet and MemBrain-Seg V2, delivering superior performance in both semantic and instance segmentation. Its ease of use, through the Napari plugin and its ability to handle diverse datasets without retraining, makes TARDIS accessible to a broader audience of researchers. With the continuous addition of new segmentation networks for various macromolecules, we envision TARDIS evolving into a comprehensive framework capable of annotating all cellular structures. By making our datasets and code publicly available, we hope to foster an open science environment that accelerates future innovations, advancing our understanding of cellular architecture and biomolecular function.

## Acknowledgments

We thank Huihui Kuang (NYU Langone Health, New York) for acquiring membrane micrographs and Fred Heberle (University of Tennessee, Knoxville) for providing membrane micrographs and testing the early version of TARDIS. We thank Aygul Ishemgulova (NYSBC, New York) for providing a tomogram in Figure 4C containing TMV structures. We thank Alex Noble (NYSBC, New York) for providing a membrane cryo-ET dataset, Misha Kopylov (NYSBC, New York) for acquiring cryo-ET microtubule dataset and Daija Bobe for (NYSBC, New York) preparing grids. We would also like to thank Neal Waxham (McGovern Medical School, Houston) for providing feedback and beta-testing the software. Special thanks to Utz Ermel, Bridget Carragher, and Clint Potter (CZI for Advanced Biological Imaging, Redwood City) for their support and help with depositing the segmentation dataset on the CZI cryo-ET data portal. We thank the SEMC IT team for technical support. The authors thank the members of the EM facility at the MPI-CBG (Dresden, Germany) for their technical support in acquiring the tomographic data of plastic-embedded samples.

This work was performed in part at the Simons Electron Microscopy Center located at the New York Structural Biology Center and was supported by grants from the Simons Foundation (SF349247) and Chan Zuckerberg Initiative (2023-331615 (5022) GB-1597298). Research in Oliver Clarke lab is supported by a grant from National Institutes of Health (1R01HL168178). Research in the Müller-Reichert lab is supported by grants from the Deutsche Forschungsgemeinschaft (DFG; MU1423/8-3, grant number 258577783 and MU1423/10-3, grant number 282354882). The Müller-Reichert lab also acknowledges support from the CCBx program of the Center for Computational Biology of the Flatiron Institute (NY, USA, SF 1157392). The research in the Shee-Mei Lok group was supported by the Ministry of Education, Singapore (MOET32023-002). Research in the Redemann lab was supported by Human Frontiers in Science HFSP (RGY0070/2019), National Institutes of Health Grant (NIH) R01GM144668 and by the CCBx program of the Center for Computational Biology of the Flatiron Institute (NY, USA, SF1157393). Magdalena Magaj was supported by the Wagner Fellowship at UVA. The work in the Bařinka lab was in part funded by the CAS (RVO: 86652036) and the Czech Science Foundation (GA23-07149S).

## Methods

### Datasets

#### Microtubule tomographic and TIRF datasets

To develop TARDIS, we assembled a comprehensive collection of tomograms obtained under both room and cryogenic temperatures (**Supp. Table 1**). This dataset includes 93 tomograms ranging in resolution from 1.16 to 3.64 nm. We then divided these datasets into training, testing, and validation subsets using a 70/20/10 split. The collection encompasses three different imaging techniques, and seven specimen preservation methods and includes samples from eight distinct species, enhancing its diversity.

Microtubule segmentation was initially performed automatically using the AmiraZIBEdition software^40^, and subsequently refined manually according to established protocols^62^. The segmentation results were saved in Amira proprietary ASCII format, detailing each microtubule instance as a set of points. For semantic segmentation, we created binary masks from point sets by drawing splines along each instance with a diameter of 25 nm^63^. This approach effectively transforms the detailed instance data into a format suitable for training and evaluating our semantic segmentation model.

We also collected a dataset from interference reflection microscopy (IRM)/TIRF microscopy. The dataset includes TIRF images and movies acquired with Nikon Ti-E microscope equipped with an H-TIRF System (Nikon, Japan), a 60× oil immersion 1.49 NA TIRF objective and an Andor iXon Ultra EMCCD camera (Andor Technology, Belfast, UK). Double-stabilized porcine microtubules were visualized using the IRM channel, while Janelia 549 labeled microtubule-binding proteins were visualized with a 561 nm excitation laser. Segmentation was performed manually in Fiji and the annotations were converted to comma-separated values (CSV) file format.

#### Membrane tomographic dataset

For our membrane dataset, we face a similar challenge to the microtubule scenario, a lack of existing datasets for tomographic reconstructions or micrographs. To address this, we assemble a diverse dataset of membranes based on expert curation. We also included tomograms featuring membrane structures from the EMPIAR and CZI Cryo-ET data portals to further broaden the dataset.

This effort results in a dataset of 53 tomograms with resolutions ranging from 0.55 to 2.40 nm. We divide the dataset into training, testing, and validation segments with a 70/20/10 split. Segmentation was carried out in two stages. First, we hand-segment 10 tomograms to train our initial FNet model, after which we use this pre-trained model to perform all following segmentation. We employed Napari with a custom interpolation plugin to curate all membranes segmentation.

Using our segmentations, we extract instances from binary masks through a connected component algorithm. Point clouds were generated via skeletonization and voxelization, ensuring a consistent and accurate representation of membrane structures in our dataset.

#### Membrane micrograph dataset

In addition to assembling a membrane dataset from tomographic reconstructions, we enhance our training and validation resources for CNN models by incorporating micrographs. This expansion involves not only using micrographs provided by collaborators but also simulating micrographs through tomogram projections. We achieve this by selecting tomographic volumes approximately 60 nm thick and applying z-max projections to replicate the appearance of micrographs. This approach significantly diversifies our dataset, allowing us to include a broader range of data types. The methods for semantic and instance segmentation of these simulated micrographs adhere to the same procedures previously outlined.

#### Actin tomographic dataset

We assemble tomograms of actin obtained under cryogenic temperatures (**Supp. Table 1**). In total we fully annotated actin in 3 tomograms with resolution 1.38 nm and 1.81 nm. We divide the dataset into training, testing, and validation segments with a 70/20/10 split.

#### Point Cloud synthetic dataset

To train the DIST model, we construct a dataset combining actual and simulated data. Real images of membranes and microtubules were hand-segmented to create ground truth masks, which are then skeletonized and voxelized. Simulated point clouds are also generated for filaments and membranes, and processed using the same methods as the real data. This approach ensures consistency between simulated and real datasets. To further augment simulated dataset, each generated point is jiggled in random direction and amplitude to imitate image noise. Additionally, simulated datasets were further augmented with random noise set-up to 10% of the number of points and random drop of points within instances of microtubules and membranes. Each point cloud is annotated with the correct instance IDs based on known ground truth positions. The dataset is divided into 80% for training and 20% for validation, with the simulated data included solely in the training set, comprising 50% of the total training dataset.

### Data preprocessing for CNN

Data preprocessing and augmentation are crucial for optimizing model performance. Initially, each micrograph and tomogram underwent intensity normalization, where the mean intensity is subtracted and divided by the standard deviation, followed by scaling all values to a range between –1 and 1. To manage large images, we cut them into overlapping patches – 256×256 pixels for micrographs and 96×96×96 voxels for tomograms. The datasets are then divided into training, testing, and validation (**Supplementary Table 1**).

### FNet CNN model for accurate semantic segmentation

The TARDIS semantic segmentation workflow relies on the FNet model, illustrated in **Supplementary Figure 1**, which features an encoder-decoder architecture with dual decoder branches, giving the network an F-like shape. The first decoder adheres to a traditional UNet design, employing skip connections to maintain spatial detail, while the secondary decoder incorporates hierarchical skip connections from all encoder levels, enhancing the model ability to capture features at multiple scales. This dual-decoder setup is finalized with a single convolution and sigmoid function for binary segmentation. Both FNet and UNet architectures include five layers each for encoding and decoding. Each layer is composed of group normalization, a dual-convolution block followed by a LeakyReLu^64^ activation function. The training was conducted using the NAdam optimizer with a learning rate of 0.0001 and binary cross-entropy (BCE) as the loss function. To standardize training between different modules and avoid overfitting, an early stopping criterion was applied, halting training if no validation loss improvement was observed over 250 epochs.

### Post-processing of semantic binary masks to point clouds

The initial segmentation produces binary masks for filaments and membranes, which include both false-positives and false-negatives (**Supp. Fig. 2A**). These inaccuracies affect the precision of subsequent instance segmentation tasks, such as those using connected component algorithms. To improve this, we convert the binary masks into point clouds through a series of steps.

First, we perform skeletonization of the binary masks to capture the core structure, followed by voxel down-sampling. The skeletonization process starts with a Euclidean distance transformation, which computes the shortest distance from each foreground pixel to the nearest background pixel, creating a distance map of the mask. We then apply thresholding based on specific feature sizes (12.5 nm for microtubules and 2 nm for membranes) to reduce noise and separate closely packed structures (**Supp. Fig. 2A**).

Next, we generate point clouds by recording the coordinates of each pixel in the skeleton. Given that these point clouds can become very dense, sometimes exceeding one million points, we use voxel down-sampling to reduce the point count by a factor of ten without losing spatial and geometric detail. This down-sampling yields a simplified yet accurate representation of the original structures (**Supp. Fig. 2B**), enhancing computational efficiency and manageability.

### DIST model for point cloud instance segmentation

Instance segmentation with DIST is divided into three parts: 1) Transformation of the raw point cloud into a pairwise representation based on the distances between points to capture the initial edge features. 2) Processing through the DIST model to output a matrix representing the probability of connection between each point pair, as shown in **Supplementary Figure 3C**. This matrix predicts potential edges in the instance graph, forming the basis for distinguishing individual instances. 3) Application of a graph cut algorithm on the maximum likelihood graph to segment instances based on the predicted edges.

Training of the DIST model focuses on predicting these edges by learning vector representations for each point pair within the cloud, termed pairwise or edge embeddings. To achieve translation and rotation invariance, initial edge features are based on the Euclidean distances between point pairs, employing a scaled exponential function of the negative squared distance to prioritize proximity. Formally, for a point cloud comprising points pi(i = 1, …, n), we construct a graph G with nodes for each pi and edges ei,j connecting all pairs, where the weight ei,j is calculated as exp(−d2i,j / (2×s2)), with di,j denoting the Euclidean distance between pi and pj, and s representing a predetermined scale factor. This weighting scheme effectively encodes the geometric relationships between all points in the cloud, leveraging Euclidean distances to capture spatial structure in a translation– and rotation-invariant manner. This allowed us to ensure the SO(n) invariance for translation and rotation and embed all geometrical information about nodes geometrical relationship and position in the space. This approach effectively addresses the challenges posed by the complex nature of objects in point clouds, especially in bioimaging domains where these objects can be large and may intersect or overlap.

### Training DIST model for instance segmentation

We train two general DIST models (**Supp. Fig 3B**). The first segments chain-like structures, for example microtubule filaments or 2D membranes derived from cryo-EM micrographs. The second, segment 3D surfaces, like membranes, mitochondria, etc. Similarly, as described before DIST was trained using the NAdam optimizer with a 0.00001 learning rate. An early stop mechanism was employed to halt training when there was no improvement in the moving average of validation loss over 100 consecutive epochs. Regarding the loss function, binary cross-entropy (BCE) was used to train the DIST model.

Training and evaluating the DIST model required building an instance graph representation for each point cloud. The graph is represented as a 2D matrix (**Supp. Fig 3A**) with nodes p represented by coordinates in n-dimensional space, and the edge represents spatial connectivity between two nodes p_i_ and p_j_, where *i* is shown in the matrix row, and *j* in the matrix column. The graph representation for linear-like structures (membranes in 2D and microtubules) is constructed from ground truth annotations in which each instance is an ordered list of nodes. Knowing the order, the edge matrix is defined as 1 for each neighboring p_i,j_. This approach imposes restrictions where each *i* can have only up to two edges. In the case of 3D structures like membranes, we build this 2D matrix by establishing for each p_i_ edge connection to its eight nearest neighbor p_j_ within the same instance.

### Graph cut from the predicted adjacency matrix

The output of the DIST model is a probability adjacency matrix (**Supp. Fig. 3A**), which contains the probability of an edge existing between a set of two points. For the segmentation of filament and membrane structures, which are known to have a linear chain we know the prior that each node (points) has at most two neighbors. We can use this information to search for the graph chain of nodes, where each node is connected to two other nodes with the highest probability. In this case, we adopt a greedy algorithm for graph inference. First, we build a hashmap for each node in the point cloud containing node ID, and edge probability p_i,j_. Next, for each new instance, we searched the hash map for the initial node. This allows us in the final step to iteratively find up to two edges with the highest probability for all connected nodes. This process is continued until no new edges can be recognized for a given instance. We use this approach when segmenting membranes and microtubules that follow this chain structure. In the case of 3D structures like membrane volumes, we do not have such priors, therefore we use a greedy graph cut algorithm that connects nodes with all other nodes if their probability of connection is higher than 0.5. This process continues until no new edges can be identified for a given instance. The output of this approach is a list of coordinates belonging to the same instance which can be output in any format suitable for the user.

### Benchmark experiments

#### Semantic segmentation

In the evaluation of the semantic segmentation capabilities of the TARDIS framework, we employed a comprehensive set of metrics, including F1 score, precision, and recall (**Eq.1-3**), alongside more nuanced measures such as average precision (AP; **Eq. 4**) and average precision at a recall threshold of 90% (AP90; **Eq. 4**). These metrics collectively offer a robust assessment of the segmentation accuracy, effectively capturing the performance of TARDIS in prioritizing true positives over false positives and negatives. The AP and AP90 scores provide insights into the model efficacy across varying thresholds of detection confidence, reflecting the area under the precision-recall curve and thus offering a detailed picture of model performance in semantic segmentation tasks.

#### Instance segmentation

For the instance segmentation evaluation, TARDIS was assessed using the mean coverage (mCov; **Eq. 5**) metric. The mCov metric is designed to accurately quantify the overlap between the ground-truth instances and the predicted instances within the point cloud, using intersection over union (IoU) as the basis of measurement. This approach ensures an accurate evaluation of instance segmentation.

#### Naïve distance grouping

In the absence of a pre-existing baseline benchmark for evaluating the performance of similar point cloud data derived from filaments and membranes, we devised our benchmarking standard, rooted in the principle of naive distance grouping. This approach entailed computing the average distance between neighboring points within each point cloud dataset. This average distance served as a critical distance threshold for each node (point) in the dataset. To delineate instances within the data, we grouped points based on the premise that any given node could be connected to a maximum of either two other nodes or an indefinite number (infinity) of nodes, provided these connections did not exceed the established threshold distance. This method allowed us to programmatically retrieve instances from complex point cloud datasets by leveraging spatial proximity.

### Phosphomimetic mutant experiments

#### U2OS NDC80c/Hec-1 mutant cell line

U2OS cells lines expressing various phosphomutant Hec1:SNAP, were generated, as described previously^48^. For electron microscopy, cells in mitosis were enriched by applying the shake-off technique^65^. Flasks with cell confluency of 60–80% were shaken against the laboratory bench. The medium with detached cells was then collected, centrifuged at 1200 rpm for 3 min at room temperature, and resuspended in 1 ml of pre-warmed DMEM medium.

#### Tomogram acquisition and microtubule dataset assembly

Cultures enriched in mitotic HeLa cells were further processed for electron microscopy essentially as described previously^10^, and cryo-immobilized using an EM ICE high-pressure freezer (Leica Microsystems, Austria). Frozen samples were then substituted with anhydrous acetone containing 1% (w/v) osmium tetroxide (EMS, USA) and 0.1% (w/v) uranyl acetate (Polysciences, USA). After freeze substitution, samples were infiltrated with pure resin and embedded.

To select cells in metaphase, resin-embedded samples were pre-inspected using an Axiolab RE upright brightfield microscope (Zeiss, Germany) with a 5x and a 40x objective lens (Zeiss, Germany). Selected cells in metaphase were sectioned using an EM UC6 ultramicrotome (Leica Microsystems, Austria), for semi-thick (∼300 nm) ribbons of serial sections. Collected samples were finally, post-stained with 2% (w/v) uranyl acetate in 70% (v/v) methanol, followed by 0.4% (w/v) lead citrate (Science Services, USA) in double-distilled water, and deposition of 20 nm-colloidal gold (British Biocell International, UK), as described previously^10^.

Serial sections were then transferred to a TECNAI F30 transmission electron microscope (Thermo Fisher Scientific, USA) operated at 300 kV and equipped with a 4k × 4k sCMOS camera (OneView, Gatan, USA). Using a dual-axis specimen holder (Type 2040, Fishione, USA), tilt series were acquired from –60° to +60° with 1° increments at a magnification of 4700 x and a final pixel size of 2.572 nm applying the SerialEM software package^66,67^.

MTs were segmented fully automatic using TARDIS. The serial tomograms of each recorded cell were stitched using the segmented MTs as alignment markers^68^, and MTs with their putative plus end associated with the chromosomes were manually defined as KMTs^10^.

## Supplement

### Example application for TARDIS

#### TARDIS annotated the entire database within a month

To demonstrate this, we segmented over 13,000 tomograms from the CZI Cryo-ET data portal enabling comparative biophysical analyses of membranes across various cell types (Fig. 5A-B), with pixel resolutions ranging from 0.867 to 56.12 Å. TARDIS autonomously processed these datasets in an average of 3.75 ±3.38 (mean ± standard deviation) minutes per tomogram, reducing the total segmentation time from an estimated 35 years (assuming one day per tomogram) to just one month.

**Figure 5:**
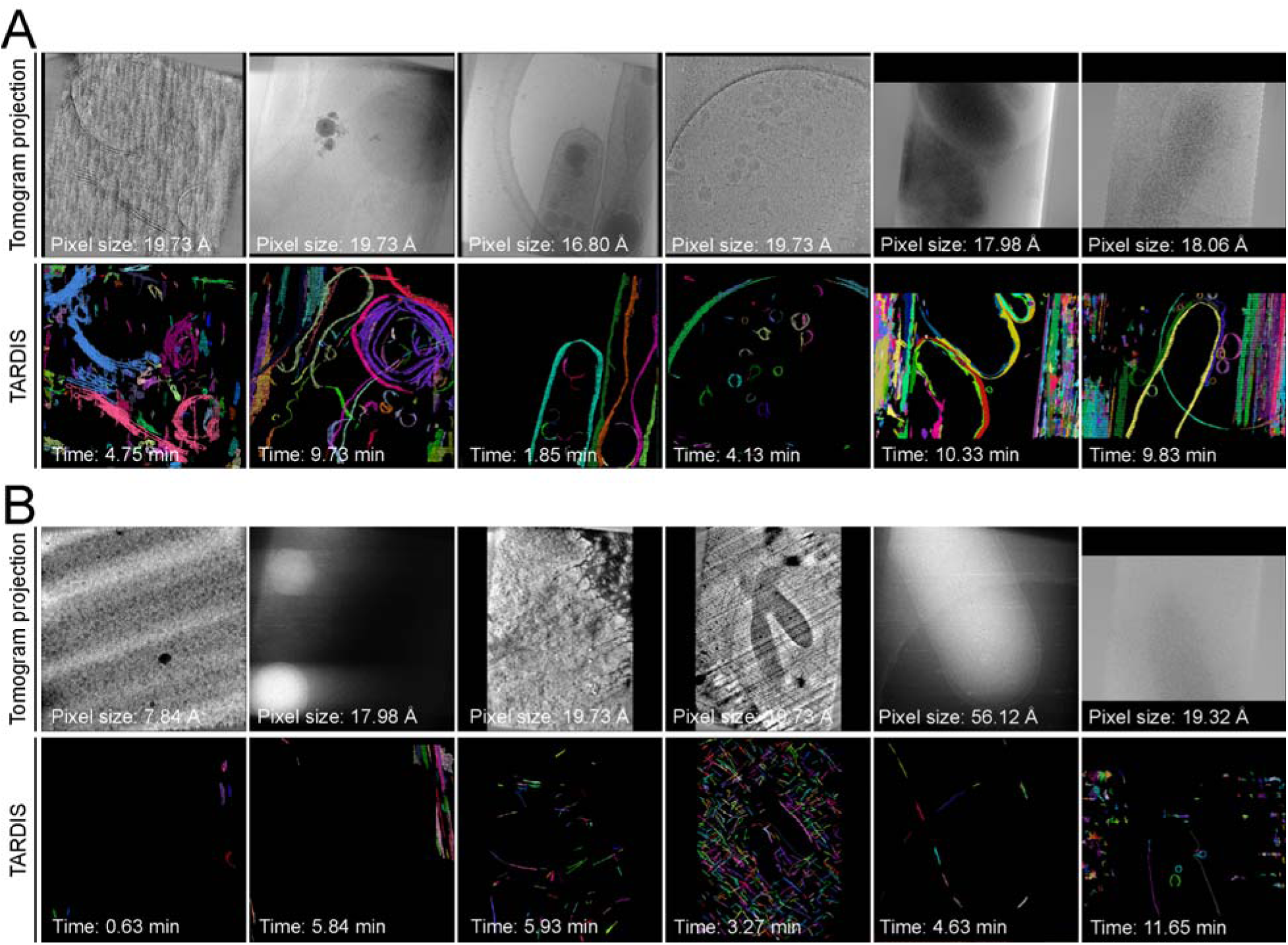
Gallery of example datasets predicted with TARDIS. **A)** Examples of TARDIS predictions on randomly selected datasets from the CZI Cryo-ET data portal, with indicated pixel sizes and segmentation times. Colors represent individual instances. **B)** Example of a TARDIS prediction on a randomly selected dataset identified by TARDIS as containing no membrane structures.

**Figure 6:**
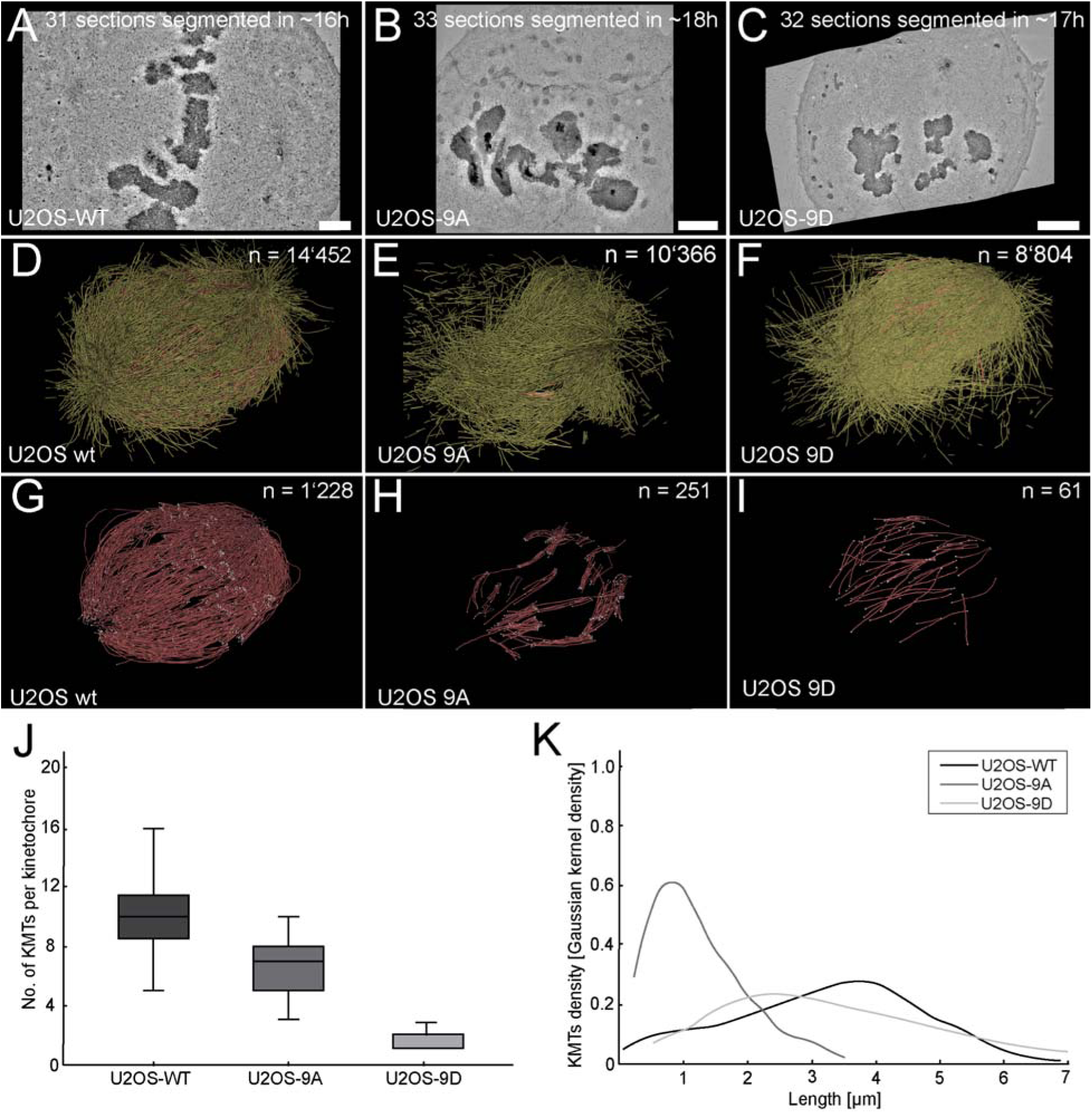
Ultrastructural reconstruction of mitotic spindles in U2OS cells from electron tomograms using TARDIS. **A)** Image of a center slice from the stitched volume of a wild-type mitotic spindle in a U2OS cell. Scale bar 1 µm. **B-C)** Center slice of a stitched volume of a U2OS phosphomimetic 9D and 9A mutant. **D-F)** 3D visualization of all MTs segmented by TARDIS. **G-I)** 3D visualization of all KMTs. **J)** Plot of the number of KMTs per kinetochore. **K)** Gaussian kernel density plot showing the KMT length distribution.

**Figure 7:**
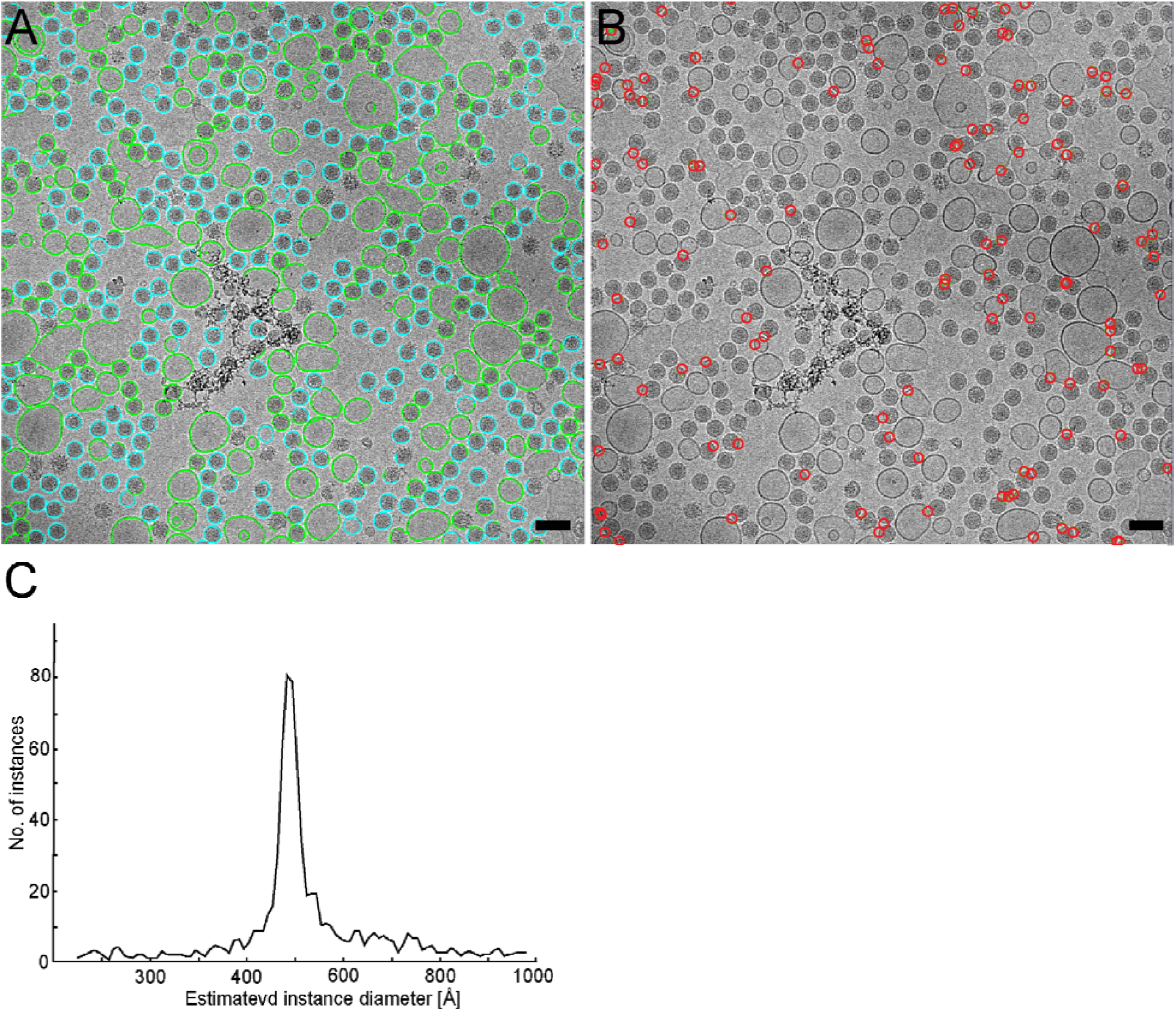
Example of TARDIS enabling biophysical study of viruses/liposome interactions. **A)** Lipid membranes segmented by TARDIS, overlaid on the corresponding micrograph, and colored as liposomes (green) or virus (blue) based on the calculated size of each segmented membrane. Scale bar 100 nm. **B)** Virus-liposome interaction sites inferred based on TARDIS segmentation in (A). **C)** Size distribution of the membrane-containing objects identified by TARDIS in the dataset shown in (B). A visible peak around 50 nm corresponds to the diameter of a mature flavivirus particle, with liposomes having a more variable size distribution.

This efficiency opens new avenues for analyzing cellular architecture. For example, automated segmentation revealed that 8.51% of tomograms lacked membrane structures or were of poor quality (**Supp. Fig. 10A-B**). The remaining tomograms contained an average of 5.59 ±7.26 (mean ± standard deviation) membranes per tomogram, with volumes of 0.009 ±0.017 µm³ (mean ± standard deviation). These large-scale quantitative insights, previously unattainable, now enable deeper exploration of cellular morphology and organization.

#### Simultaneous analysis of membrane curvature and interaction with microtubules using TARDIS

To further demonstrate TARDIS capabilities, we segmented 75 tomograms from the DS-10160 and DS-10161 CZI datasets^18^ of the bacterium *Hylemonella gracilis* under varying conditions, completing the task in ∼5 hours on a single A100 GPU. This analysis revealed inner and outer membrane curvatures of 0.026 and 0.019, respectively. Exposure to Bdellovibrio shifted these values to 0.023 and 0.021, suggesting membrane softening or remodeling in response to the predator^69^ (**Supp. Fig. 11A-B**).

#### TARDIS uncover microtubule/membrane interaction in mouse axones

We applied TARDIS to 30 tomograms from the EMPIAR-10815 dataset^70^, focusing on axons from mouse dorsal root ganglia. In ∼2 hours, we segmented both membranes and microtubules, identifying an average of 4.8 ±2.5 (mean ± standard deviation) microtubules per field of view (**Supp. Fig. 11C**). Notably, 3 microtubules per field were consistently located within 50 nm of a membrane, with an average length of 173 nm, indicating potential microtubule-membrane interactions that may support anchoring and transport functions.

We tried to replicate this experiment both with MemBrain-seg V2 and Amira respectively for membrane and microtubule segmentation. This however was unsuccessful, as MemBrain-seg was not able to accurately segment individual instances (**Supp. Fig. 11A-B**), and in case of microtubules, Amira could not output even single microtubules. This one more time presents the issue with the state of currently available tools which do not allow for streamline processing of new, unseen dataset requiring extensive fine-tuning training or fallback to manual annotation.

## Ablation study

### TARDIS improves semantic segmentation performance due to the FNet architecture

TARDIS incorporates a novel CNN architecture called FNet into its workflow. We evaluated FNet on semantic segmentation tasks involving microtubules and membranes and found that the TARDIS FNet model significantly outperforms current state-of-the-art tools (**Table 1-3**). To compare TARDIS FNet efficiency also against other CNN architecture, we trained both FNet and a similarly deep UNet model within TARDIS, ensuring identical training parameters for a fair comparison. As expected, the TARDIS FNet model achieved superior results compared to the TARDIS UNet model across both microtubule and membrane benchmarks (**Suppl. Fig. 7A-C**). Specifically, the FNet model attained better metrics for microtubule segmentation from both cryo-ET and plastic ET datasets (**Suppl. Fig. 7A, Supp. Fig 8A** and **Suppl. Table 5**). Additionally, TARDIS FNet demonstrated significant performance improvements in membrane segmentation tasks using data from both cryo-ET (**Suppl. Fig. 7B, Supp. Fig 8B** and **Suppl. Table 6**) and micrographs datasets (**Suppl. Fig. 7C, Supp. Fig 8C** and **Suppl. Table 7**). Overall, the TARDIS FNet model demonstrated greater performance compared to a similarly deep UNet model. This improvement was achieved through our dual-decoder strategy, which enabled the generation of sharper probability maps (**Suppl. Fig 9**)

To further increase the reliability of TARDIS semantic segmentation, we implemented a pixel size normalization strategy. This approach ensures that various objects segmented by our TARDIS semantic segmentation module maintain at a consistent scale. We determined the optimal pixel size scaling for membrane segmentation tasks in both tomograms and micrographs by training the TARDIS FNet and UNet models with pixel normalization set to 15 Å, 8 Å, and 4 Å using micrograph datasets. Our results indicated that for tomographic datasets, the TARDIS FNet model performed best when tomograms were scaled to a resolution of 15 Å (**Suppl. Fig. 8D** and **Suppl. Table 6**). In the case of micrographs, optimal performance was achieved when the models were trained on data normalized to a resolution of 4 Å (**Suppl. Fig. 8C** and **Suppl. Table 7**). Additionally, we observed that the TARDIS FNet model set to 4 Å yielded the best results particularly for high-resolution micrographs (<10 Å), while for low-resolution micrographs, scaling to 8 Å provided the most accurate segmentation. Based on these findings, we have integrated the optimal pixel size normalization into our automated workflow and publicly released the enhanced model.

### DIST can segment other point cloud data

DIST was initially developed for cryo-electron microscopy data, but it can be also effectively applied to point cloud segmentation in domains like LiDAR from the ScanNetV2 dataset or the PartNet dataset. To test this, we applied the DIST model, specifically pre-trained on the LiDAR dataset, to segment point clouds from the ScanNetV2 and PartNet datasets. Our results demonstrate that DIST performs well on this task, effectively segmenting objects and parts (**Suppl. Fig 9D**). This shows that the DIST model generalizes well across datasets and adapt for diverse point cloud segmentation challenges, further validating its potential for tasks beyond its original bioimaging application.

**Supplementary Figure 1:**
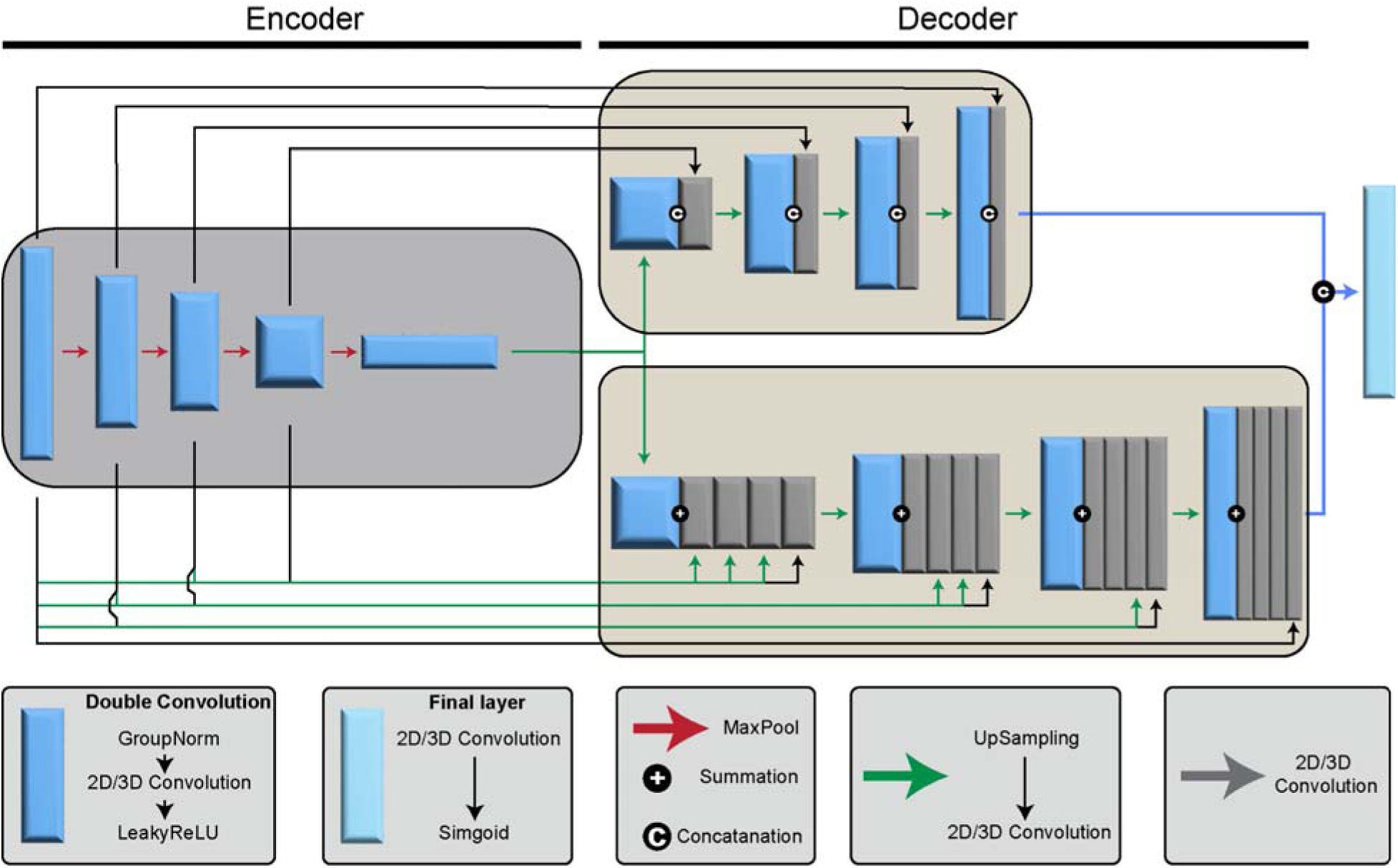
FNet CNN model architecture. The figure illustrates a CNN encoder-decoder framework dedicated to semantic segmentation in images with low signal-to-noise ratios.

**Supplementary Figure 2:**
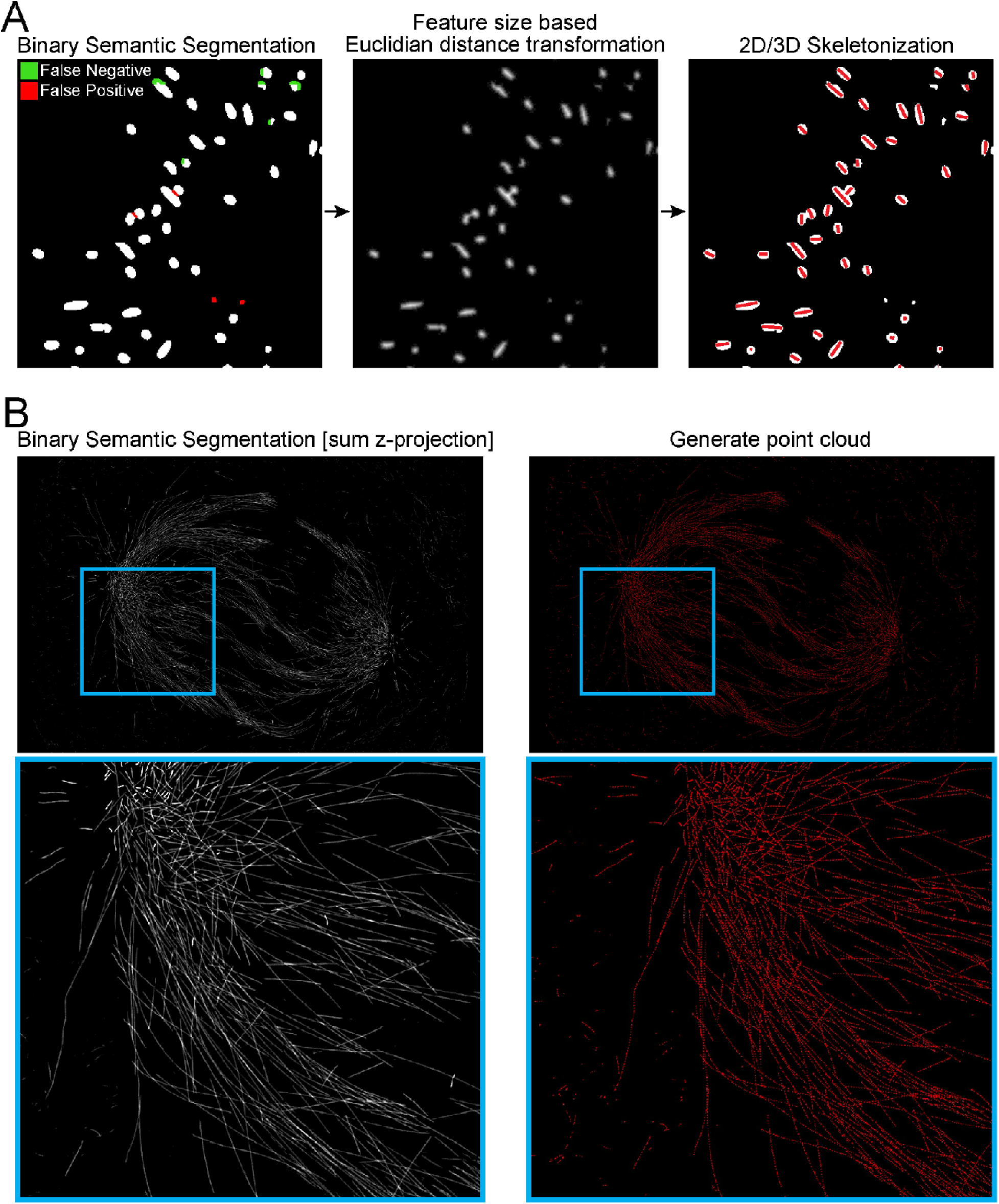
Illustration of pre-processing steps after the semantic segmentation. **A)** Illustration of converting binary semantic masks to a point cloud. Green highlights false negatives and red indicates false positives. The process begins with applying Euclidean distance transformation based on feature size to minimize segmentation errors. This is followed by skeletonization to extract the core features of the mask, which are then represented as a point cloud. **B)** Example of TARDIS binary semantic segmentation mask and the corresponding point cloud. The gray color represents a z-sum projection of the semantic mask, while the red color denotes individual points within the generated point cloud.

**Supplementary Figure 3:**
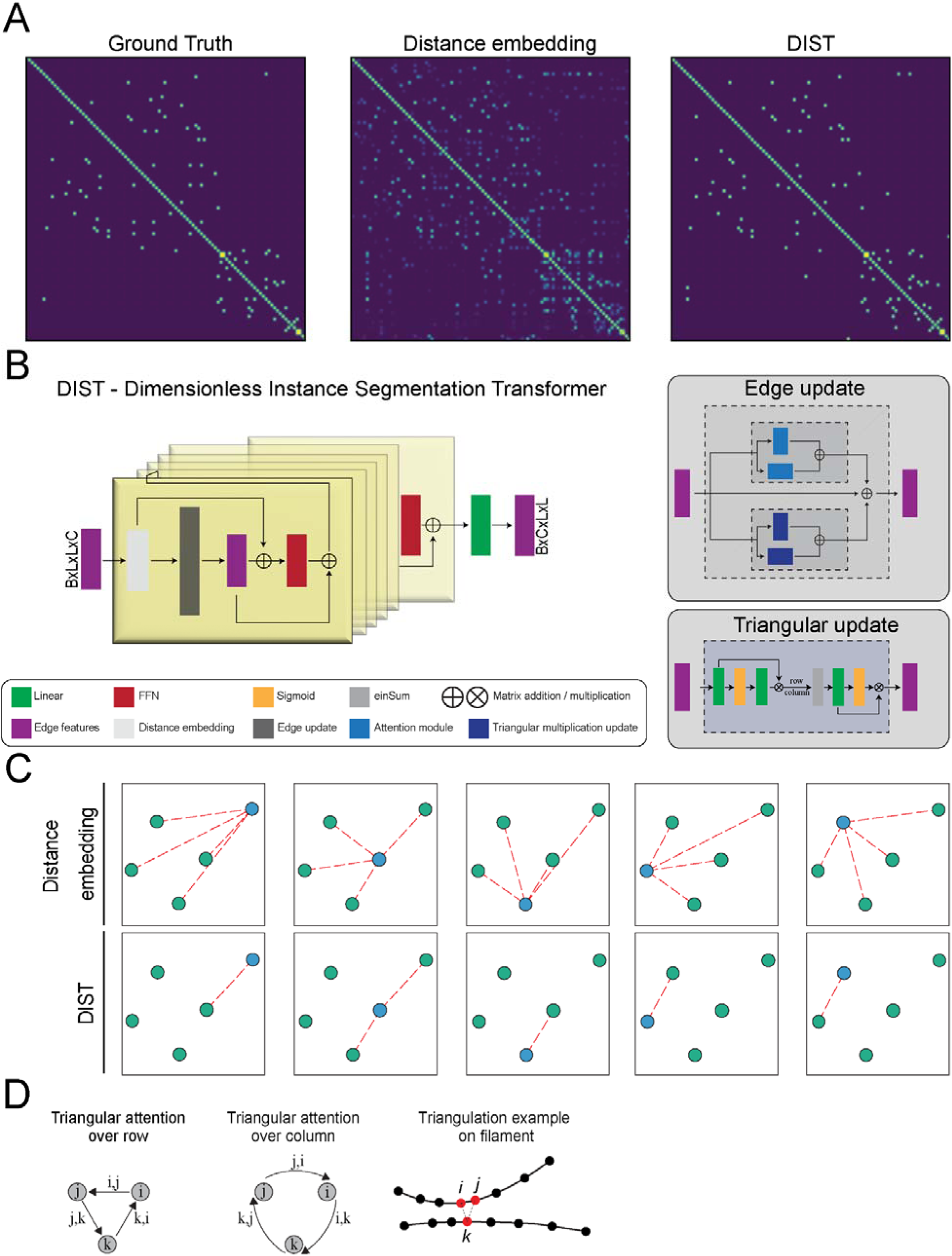
The DIST model architecture. **A)** Adjacency graphs illustrating edge connectivity in point clouds. The first graph displays the ground truth adjacency matrix with each node connected to its two nearest neighbors. The second graph shows the adjacency matrix derived from node distances and predicted probabilities using DIST. **B)** Detailed workflow of DIST starts with edge representations initialization based on distances between points in the input point cloud, resulting in an SO(n) invariant representation. DIST layers then refine edge features through triangular multiplicative updates and axial attention modules^71^, ultimately decoding these into edge probabilities. Arrows indicate the flow of information throughout the process. **C)** A cartoon depicting the distinction between initial distance embeddings and DIST predictions. Initially, each node connects to all other nodes, whereas DIST predictions retain only the relevant edges. **D)** Illustration of a triangular multiplicative update. Pairs of nodes are connected as edges on the graph, with the update process demonstrated by a triangle multiplicative update. Circles represent nodes, and arrows indicate the edges involved in the update.

**Supplementary Figure 4:**
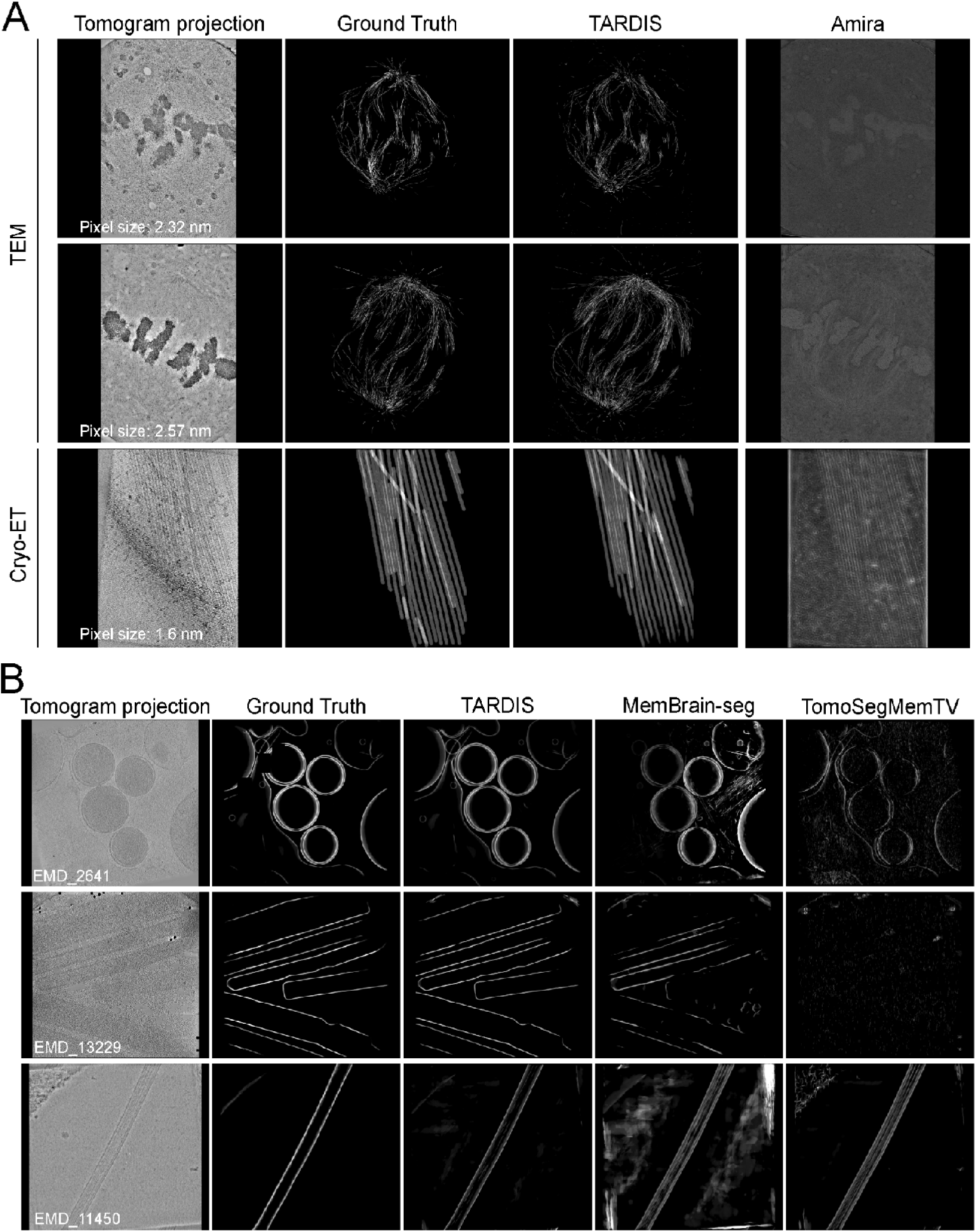
Comparison of semantic segmentations using pre-trained FNet and UNet. **A)** Predicted semantic segmentation results for tomograms containing microtubules. **B)** Predicted semantic segmentation results for tomograms containing membranes, with the gray color representing a sum z-projection of the semantic mask.

**Supplementary Figure 5:**
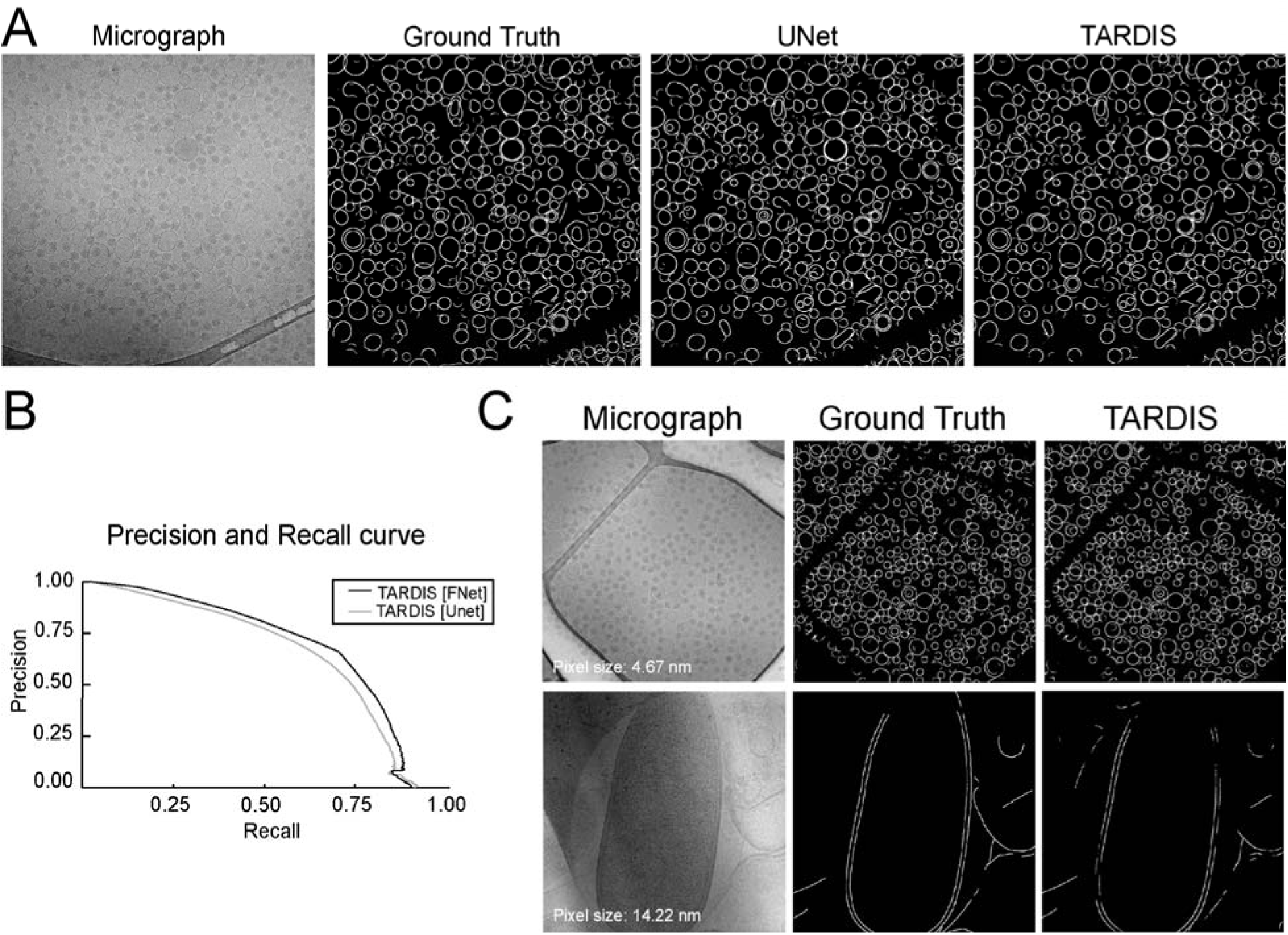
Comparison of semantic segmentations of membranes from EM micrographs. **A)** Predicted semantic membrane segmentation results for micrographs containing enveloped virus and liposome particles using TARDIS FNet and UNet pre-trained models. **B)** Precision/Recall curves for microtubule segmentation benchmark datasets, comparing predictions made by Amira, the pre-trained UNet, and TARDIS. **C)** Predicted semantic segmentation results for micrographs containing virus and liposomal membranes, on high– and low-resolution.

**Supplementary Figure 6:**
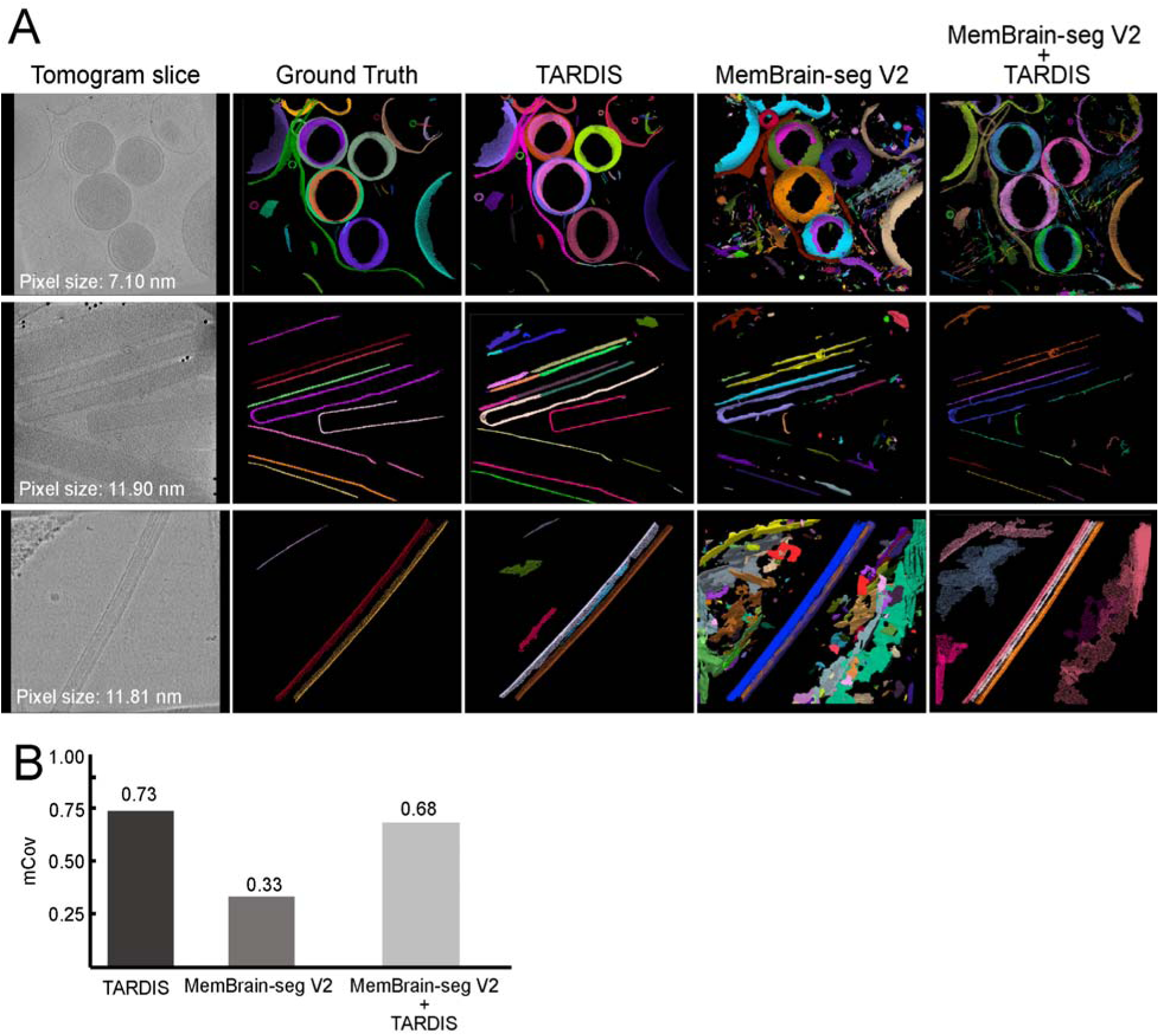
Comparison of results combining TARDIS workflow MemBrain-seg V2. **A)** Sum z-projections of predicted instance segmentations from each tomogram, with randomly assigned colors representing individual instances. **B)** Plot displaying mCov scores for instance segmentation predictions.

**Supplementary Figure 7:**
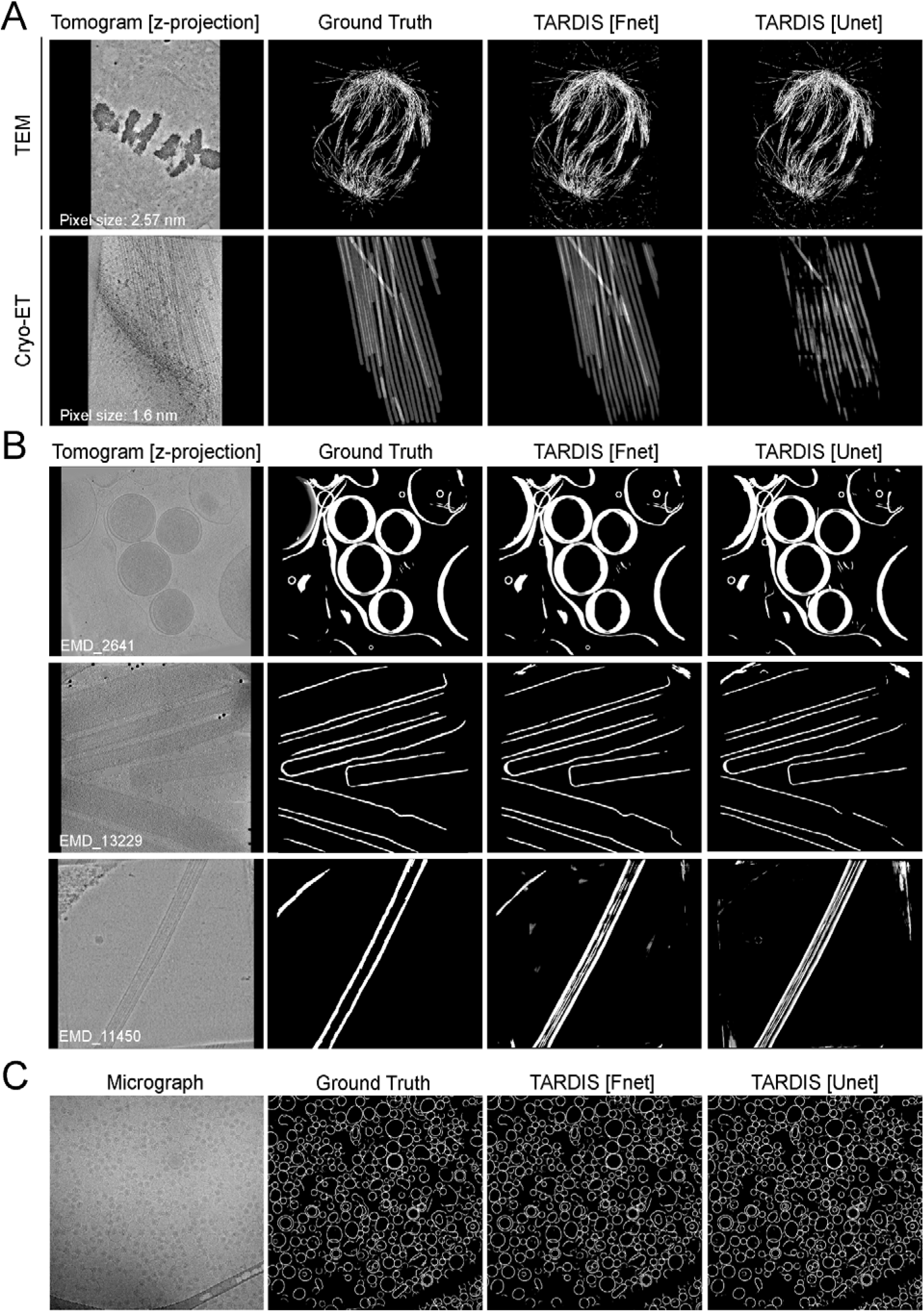
Comparison of semantic segmentations between TARDIS FNet CNN model and UNet. **A)** Predicted semantic segmentation results for tomograms containing microtubules using TARDIS FNet and UNet pre-trained models. **B)** Predicted semantic segmentation results for tomograms containing membranes. **C)** Predicted semantic membrane segmentation results for micrographs containing enveloped virus and liposomes particles.

**Supplementary Figure 8:**
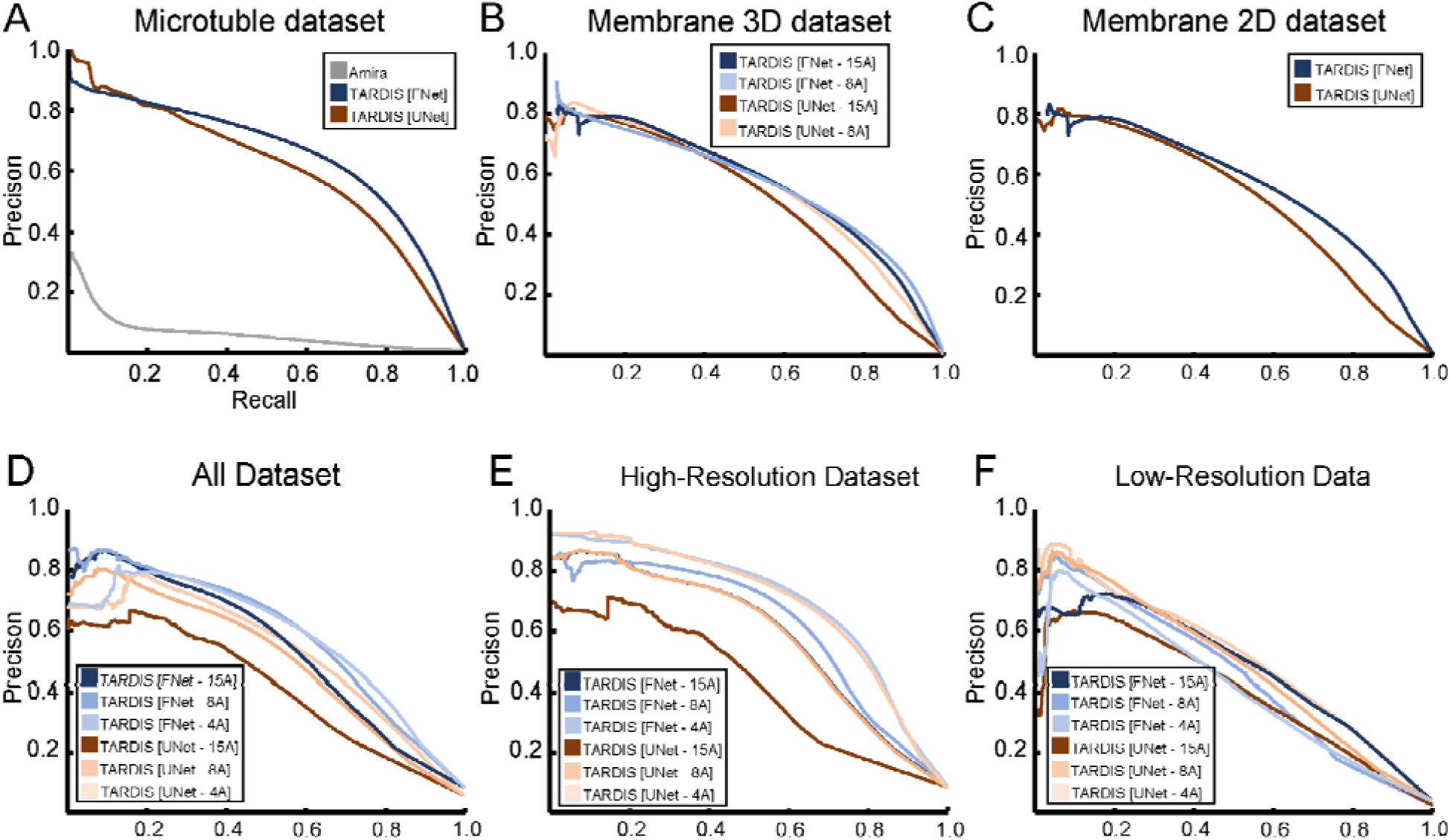
Precision/Recall curves comparing semantic segmentation performance. **A)** Precision/Recall curves for microtubule segmentation benchmark datasets, comparing predictions made by Amira, the pre-trained UNet, and TARDIS. **B)** Precision/Recall curves for membrane segmentation benchmark datasets using the pre-trained FNet and UNet model on tomographic data. **C)** Precision/Recall curves for membrane segmentation benchmark datasets using TARDIS and the pre-trained UNet model on micrograph data. **D)** Precision and recall curves for membrane datasets benchmarked using the FNet and UNet model. Different lines represent UNet models trained with specific normalized pixel size resolutions. **E)** Precision and recall curves for high-resolution membrane dataset benchmarks using the FNet and UNet model. Different lines represent UNet models trained with various normalized pixel size resolutions. **F)** Precision and recall curves for low-resolution membrane dataset benchmarks using the FNet and UNet model. Different lines represent UNet models trained with various normalized pixel size resolutions.

**Supplementary Figure 9:**
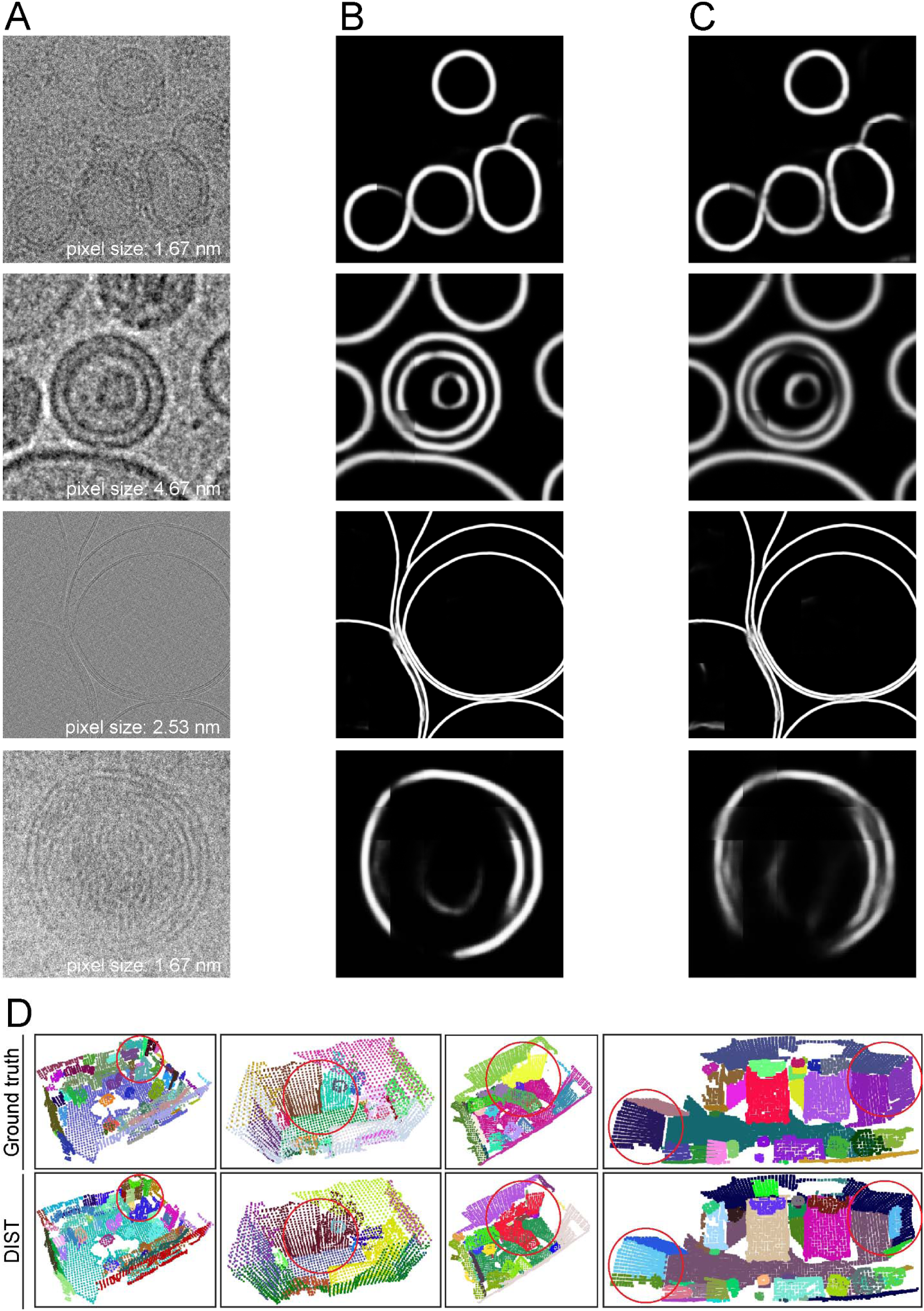
Example of Unet and Fnet prediction on micrograph with membranes at various resolutions. **A)** Cropped micrographs showing selected areas containing membranes. **B)** Probability map generated by TARDIS semantic segmentation for the area depicted in A. **C)** Probability maps of the same areas shown in A, as predicted by the UNet model. **D)** Example of point clouds segmented with TARDIS using pre-trained model on LiDAR ScanNet V2 dataset. Read circles highlight areas of discrepancy between ground truth labels and DIST predictions.

**Supplementary Figure 10:**
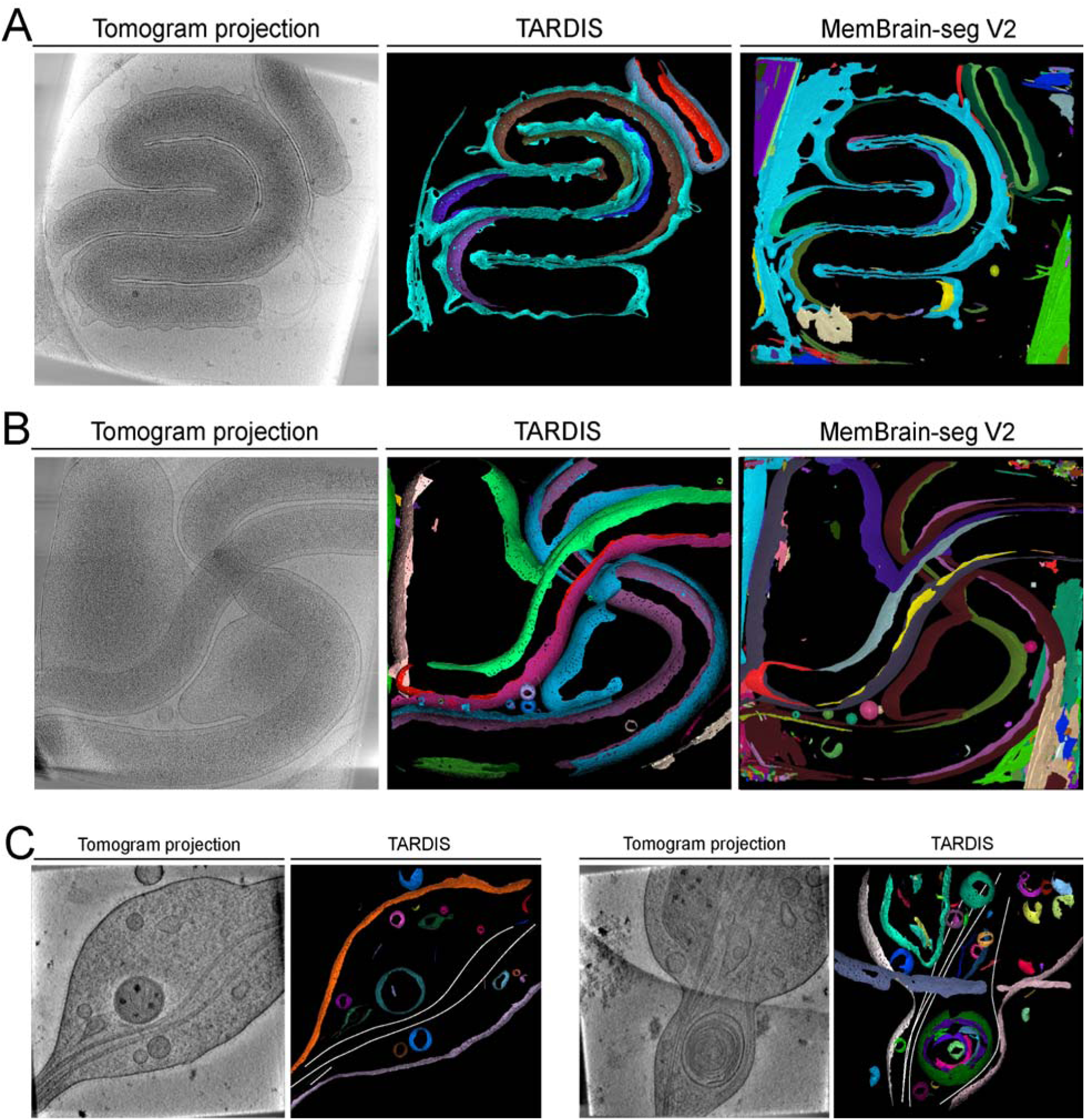
Example of TARDIS fully automated segmentation of membranes and microtubules. **A)** Example of tomographic slice and 3D view of segmented membraned from *Hylemonella gracilis*. **B)** Example of tomographic slice and 3D view of segmented membraned from *Hylemonella gracilis* in a presence of *Bdellovibrio*. **C)** Example of TARDIS segmentation of microtubules and membrane from EMPIAR-10815 dataset.

**Supplementary Table 1:**
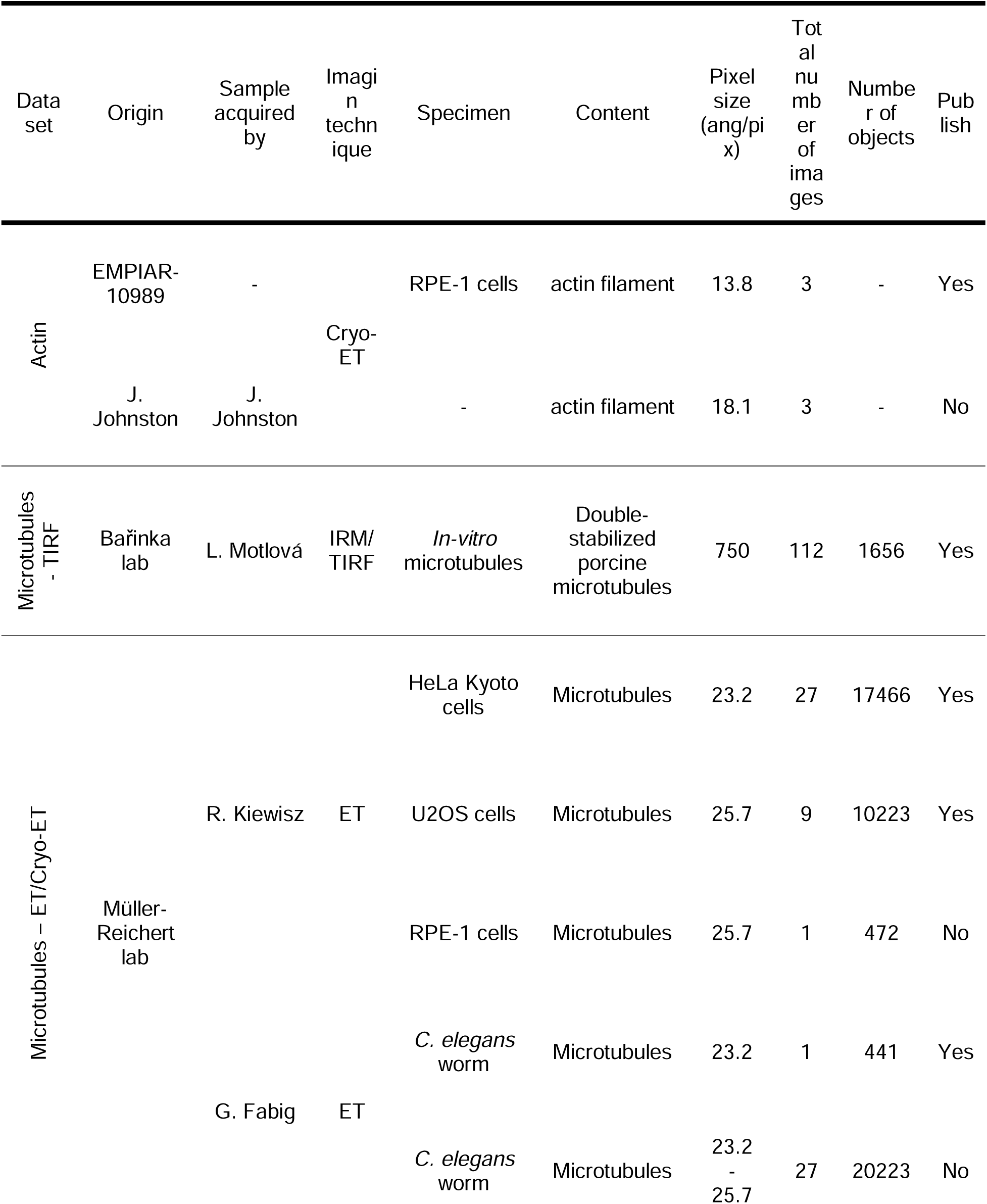
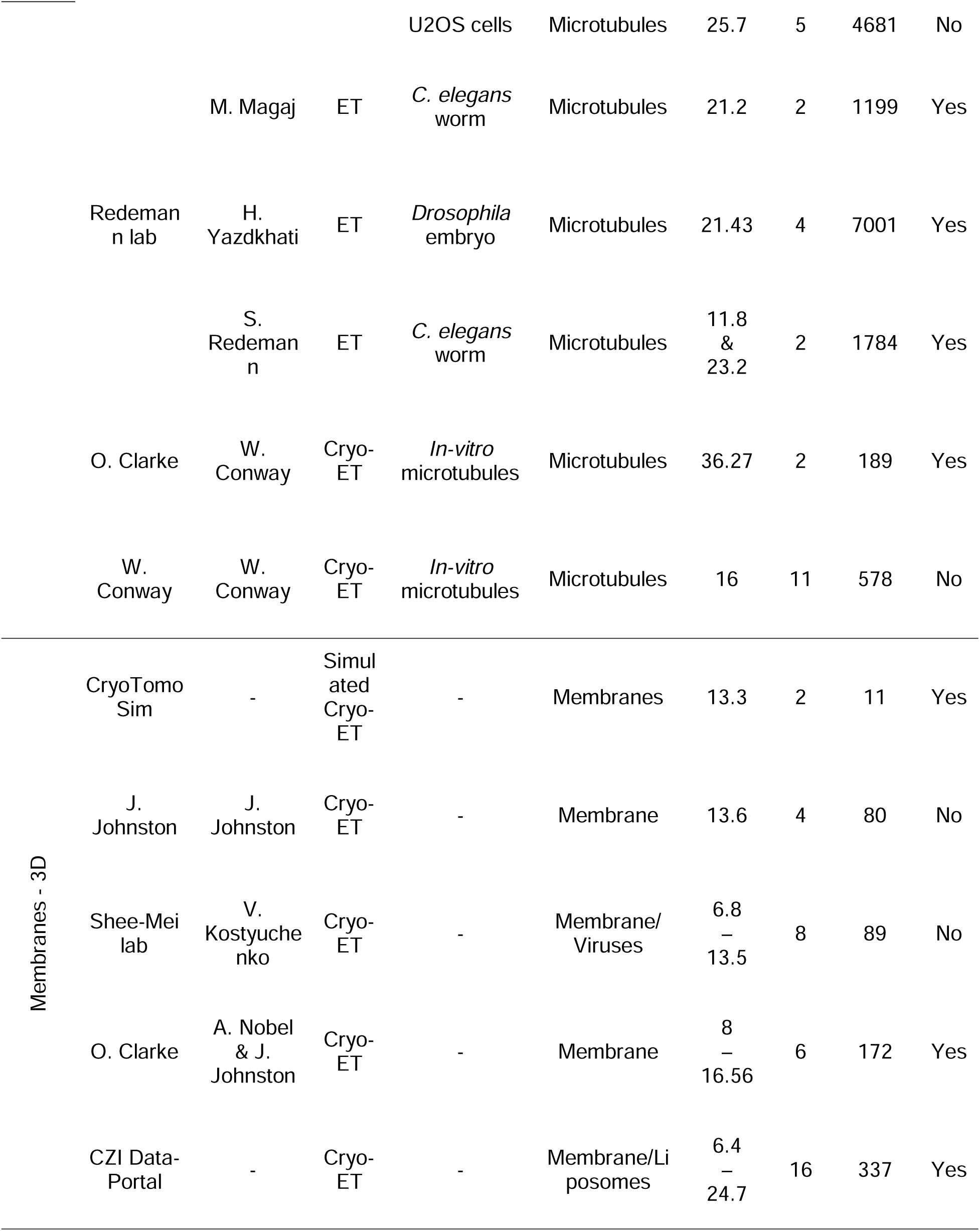
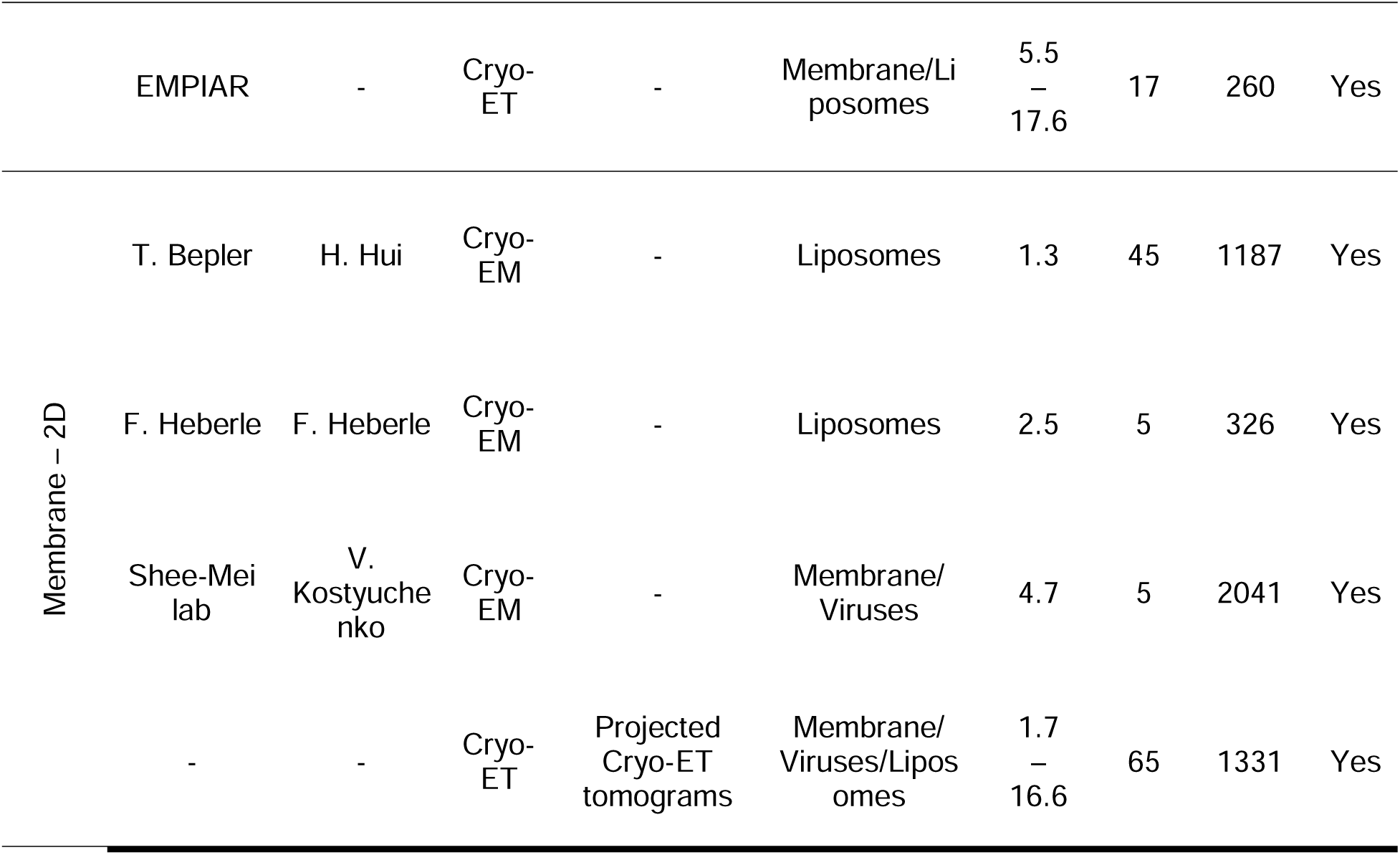
List of datasets with indicated train/validation/evaluation split.

**Supplementary Table 2:** Validation of a point cloud segmentation on ground truth data using DIST.

|  | **Filament-like object** | **Surface structures** |
| --- | --- | --- |
| DIST (linear / surface) | **0.96** | **0.91** |
| Distance grouping | 0.21 | 0.25 |

**Supplementary Table 3:**
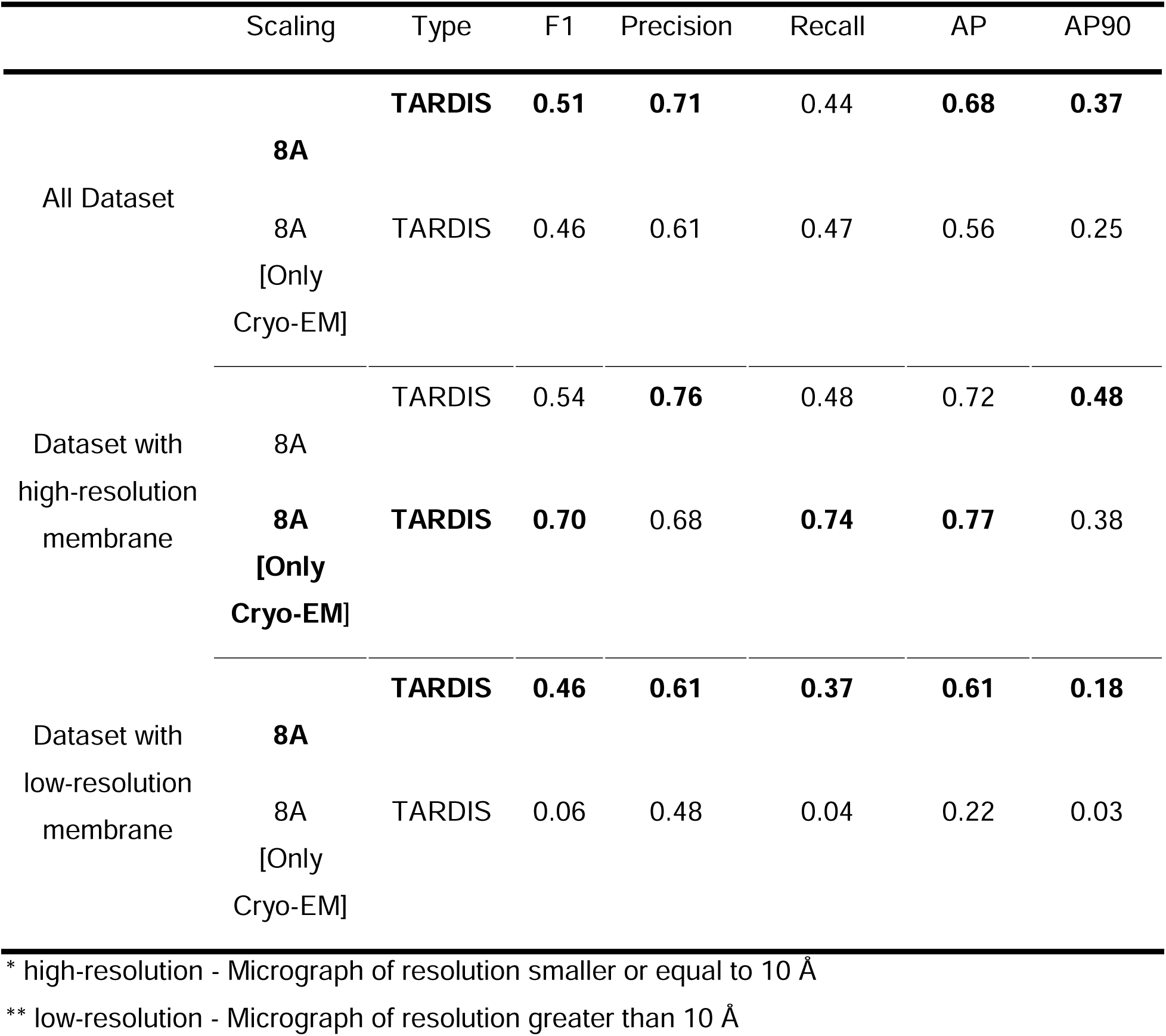
Validation of an influence of cryo-ET projection on improving semantic segmentation of cryo-EM data.

**Supplementary Table 4:** Validation of semantic and instance segmentation prediction of actin from cryo-ET tomograms.

| Type | F1 | Precision | Recall | AP | AP90 | mCov |
| --- | --- | --- | --- | --- | --- | --- |
| **FNet** | **0.07** | **0.58** | **0.04** | **0.54** | **0.06** | **0.12** |
| UNet | 0.06 | 0.55 | 0.04 | 0.50 | 0.02 | 0.10 |

**Supplementary Table 5:** Validation of semantic segmentation prediction of microtubule filaments segmented from tomographic dataset. Precision, recall, and F1 scores are calculated using a probability threshold of 0.5 for TARDIS and Amira.

|  | Model | F1 | Precision | Recall | AP | AP90 |
| --- | --- | --- | --- | --- | --- | --- |
| All<br>Datasets | **TARDIS [FNet]** | **0.61** | **0.70** | **0.55** | **0.69** | **0.33** |
|  | TARDIS [UNet] | 0.60 | 0.69 | 0.55 | 0.27 | 0.31 |
| Cryo-ET<br>dataset | **TARDIS [FNet]** | **0.58** | **0.72** | 0.48 | **0.71** | 0.20 |
|  | TARDIS [UNet] | 0.54 | 0.58 | **0.50** | 0.70 | **0.30** |
| TEM<br>dataset | **TARDIS [FNet]** | **0.61** | **0.70** | **0.56** | **0.67** | 0.35 |
|  | TARDIS [UNet] | 0.61 | 0.70 | 0.55 | 0.65 | **0.37** |

**Supplementary Table 6:**
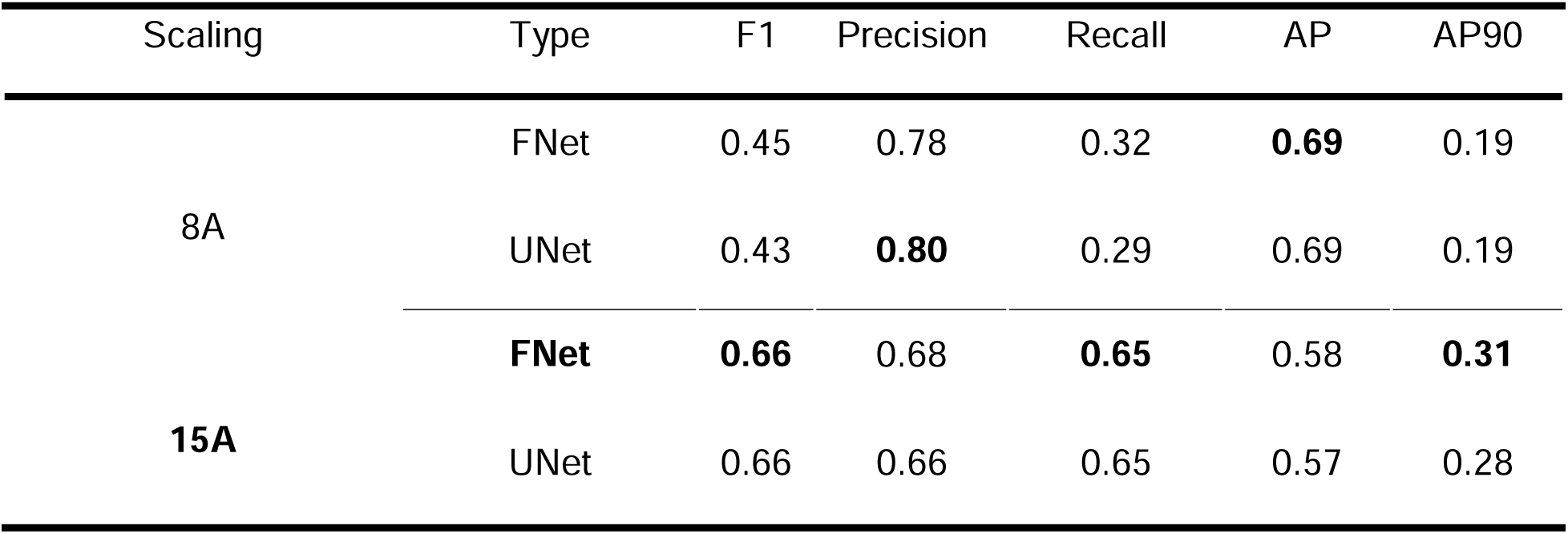
Validation of pixel size normalization on semantic segmentation prediction of membranes from cryo-ET tomograms.

**Supplementary Table 7:** Validation of pixel size normalization on semantic segmentation prediction of membranes from cryo-EM micrographs.

|  | Scaling | Type | F1 | Precision | Recall | AP | AP90 |
| --- | --- | --- | --- | --- | --- | --- | --- |
| All Dataset | 4A | FNet | **0.56** | 0.62 | **0.55** | 0.61 | **0.37** |
|  |  | UNet | 0.54 | 0.64 | 0.51 | 0.61 | 0.37 |
|  | 8A | FNet | 0.51 | **0.71** | 0.44 | **0.68** | 0.37 |
|  |  | UNet | 0.49 | 0.67 | 0.43 | 0.64 | 0.34 |
|  | 15A | FNet | 0.46 | 0.68 | 0.41 | 0.66 | 0.33 |
|  |  | UNet | 0.38 | 0.63 | 0.31 | 0.61 | 0.26 |
| Dataset with high-resolution membrane | 4A | FNet | **0.64** | 0.73 | **0.62** | 0.71 | **0.52** |
|  |  | UNet | 0.62 | 0.74 | 0.58 | 0.71 | 0.52 |
|  | 8A | FNet | 0.54 | **0.76** | 0.48 | **0.72** | 0.48 |
|  |  | UNet | 0.52 | 0.70 | 0.48 | 0.67 | 0.48 |
|  | 15A | FNet | 0.47 | 0.73 | 0.44 | 0.69 | 0.44 |
|  |  | UNet | 0.35 | 0.62 | 0.28 | 0.59 | 0.30 |
| Dataset with low-resolution membrane | 4A | FNet | 0.41 | 0.44 | **0.42** | 0.44 | 0.11 |
|  |  | UNet | 0.40 | 0.47 | 0.37 | 0.45 | 0.10 |
|  | 8A | FNet | **0.46** | 0.61 | 0.37 | 0.61 | 0.18 |
|  |  | UNet | 0.44 | 0.60 | 0.35 | 0.59 | 0.14 |
|  | 15A | FNet | 0.44 | 0.60 | 0.35 | 0.59 | 0.14 |
|  |  | UNet | 0.44 | **0.64** | 0.35 | **0.63** | **0.20** |

\* high-resolution - Micrograph of resolution smaller or equal to 10 Å
\*\* low-resolution - Micrograph of resolution greater than 10 Å

## Algorithms, Program Codes, and Listings

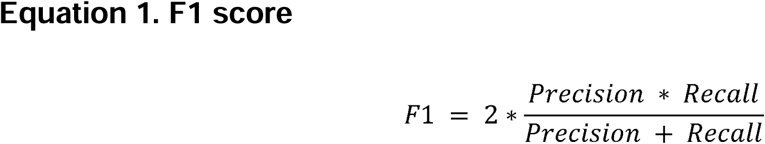

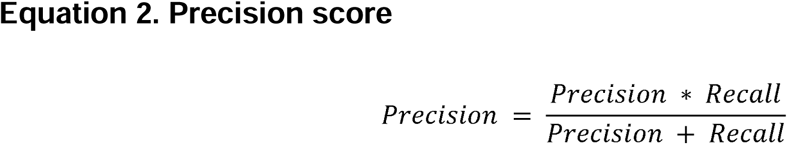

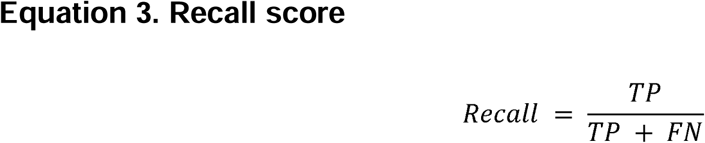

Where TP denotes the number of true positive predictions, and FN denotes false negatives.

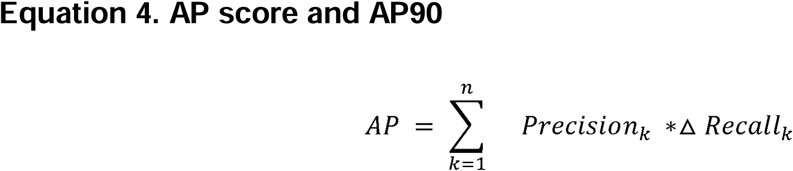

Where n is the number of thresholds, and k is the threshold value between 0 and 1. The Precision_k_ denotes precision at threshold k and ΔRecall_k_ denotes a change in recall between k and k-1.

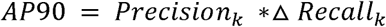

Where k denotes Precision at the threshold where recall is equal to 90%, and ΔRecall_k_ denotes a change in recall between k and k-1.

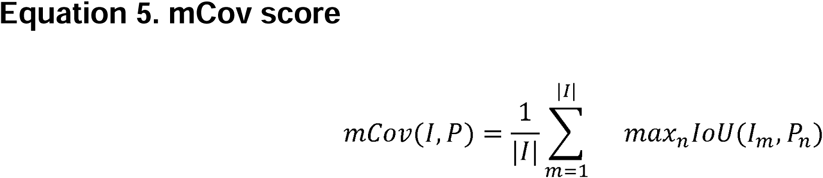

Where I_m_ denotes the number of projected spline or volume for the m-th ground truth instance. P_m_ represents the n-th predicted instance, and |I| is the number of all ground truth instances.

## Notes

https://github.com/SMLC-NYSBC/TARDIS

https://github.com/SMLC-NYSBC/napari-tardis_em

### Competing Interest Statement

The authors have declared no competing interest.

### Summary of Updates

We revise text of the manuscript to better reflect how much TARDIS is a unique approach for instance segmentation from any electron microscopy 2D/3D micrographs. We also include 2 novel study that were enable by TARDIS and were achieved in a significantly shorter time compare to other workflows.

